# Low testosterone promotes anxiety through astrocytic mitochondrial remodelling at the nucleus accumbens blood-brain barrier

**DOI:** 10.64898/2026.09.03.749088

**Authors:** Haissa de Castro Abrantes, Lewis Depaauw-Holt, Dogukan H. Ulgen, Camilla Di Giulio, Margherita Barbetti, Elias Gebara, Fiona Hollis, José L. Solano, Luise Schlotterose, Caroline Menard, Mootaz Salman, Simone Astori, Carmen Sandi

## Abstract

Low testosterone is linked to anxiety in men, but its causal role and underlying brain mechanisms remain unknown. Here, using outbred male rats stratified by natural variation in anxiety-like behaviour, we establish its causal endocrine contribution and uncover an astrocytic mitochondrial mechanism at the nucleus accumbens (NAc) blood-brain barrier (BBB). High-anxiety rats showed a NAc-selective increase in BBB permeability, alongside reduced astrocytic endfoot coverage, disrupted endfoot mitochondrial organisation and mitochondria-endoplasmic reticulum contacts, and lower mitofusin 2 (Mfn2). Testosterone suppression increased anxiety in low-anxiety rats, whereas physiological testosterone restoration reduced anxiety, reconfigured astrocytic mitochondrial and BBB-related transcriptional programmes, increased astrocytic endfoot coverage and Mfn2, recovering BBB permeability, with related effects in aged low-testosterone rats. NAc androgen receptor (AR) knockdown blunted behavioural and neurovascular responses, whereas NAc astrocyte-specific Mfn2 overexpression reduced anxiety. Thus, testosterone-AR signalling engages an astrocytic mitochondrial programme at the NAc neurovascular interface linking low testosterone to anxiety vulnerability.

## Main

Persistent high-anxiety is a major risk factor for stress-related psychopathologies and impaired well-being^1–3^, yet the biological mechanisms that sustain this vulnerability remain incompletely understood. Current frameworks have focused largely on neural circuits^4–8^ and genetic risk^9,10^, yielding major insights into anxiety vulnerability. However, affective traits are also shaped by systemic physiology^11,12^, and much less is known about how peripheral physiology contributes to brain mechanisms that sustain trait-like anxiety.

One endocrine signal long implicated in anxiety-related phenotypes is testosterone. Testosterone, present in both sexes, is the predominant circulating androgen in males, in whom levels vary with age, metabolic status, illness, and stress exposure^13–16^. In men, lower testosterone or reduced androgen signalling has been associated with psychological and anxiety-related symptoms in community cohorts, among individuals with clinical hypogonadism and during androgen-deprivation therapy^17–22^. Understanding this relationship is particularly relevant in light of reports of declining testosterone concentrations in men over recent decades^23–25^, including among adolescents and young adults^26^. Low testosterone also occurs with ageing^13,14,27^ and in the context of stress^28,29^, metabolic dysfunction^30^ and certain treatments^31^. Although anxiety phenotypes are heterogeneous and cannot be reduced to a single endocrine state, these observations identify low testosterone as a plausible, biologically tractable contributor to persistent high-anxiety in males.

Experimental manipulation of testosterone levels or androgen receptor (AR) signalling alters anxiety-like behaviour in rodents, supporting a causal role for androgen signalling in regulating these behaviours^32–36^. However, this work has largely relied on gonadectomy, exogenous testosterone administration or targeted AR manipulation. Although these approaches establish that androgen signalling can modulate anxiety-like behaviour, they do not determine whether endogenous low testosterone contributes causally to trait-like anxiety, or identify the brain mechanisms that link circulating androgen state to behavioural vulnerability.

The blood-brain barrier (BBB) is a plausible site at which androgen state could influence brain homeostasis. Androgen signalling influences endothelial function and vascular tone^15,37^, but research has largely focused on the vascular dysfunction associated with supraphysiological testosterone or anabolic-androgenic steroid exposure^38,39^. By contrast, whether reduced androgen action compromises the neurovascular interface that maintains brain homeostasis remains poorly understood. Formed by specialised endothelial cells joined by tight junctions and supported by pericytes, basement membrane components and astrocytic endfeet, the BBB regulates exchange between blood and brain parenchyma and preserves the extracellular environment required for neural signalling^40–43^. Rodent models indicate that androgen depletion or impaired neural AR signalling can increase BBB permeability and disrupt tight-junction organisation, whereas testosterone restoration can normalise barrier integrity^44–46^. In parallel, stress models implicate BBB dysfunction in affective vulnerability. Chronic social stress compromises BBB integrity in susceptible, but not resilient, animals, thereby promoting depression-related behaviours^47,48^. Stress-induced BBB alterations are also region- and sex-specific across mood-related circuits^47–49^. Together, these observations indicate that androgen state can influence BBB properties and that BBB dysfunction can contribute to affective vulnerability. Whether low testosterone contributes to regionally selective BBB alterations associated with naturally occurring differences in anxiety remains unknown.

Here we use an outbred rat model of natural variation in anxiety-like behaviour to test whether low testosterone contributes causally to a persistent high-anxiety phenotype and whether this involves a regionally selective neurovascular mechanism. We show that high-anxiety male rats have reduced circulating testosterone and that pharmacological suppression of testosterone in low-anxiety rats increases anxiety-like behaviour. We then map BBB permeability across candidate brain regions and identify increased permeability selectively in the nucleus accumbens (NAc), where it correlates with anxiety-like behaviour. We further show that testosterone supplementation shifts key NAc neurovascular features towards those observed in low-anxiety rats and reduces anxiety-like behaviours in both young adult high-anxiety rats and older low-testosterone male rats. NAc AR knockdown attenuates the behavioural and neurovascular effects of testosterone supplementation. Finally, we identify astrocytic mitochondrial remodelling as a component of this testosterone-sensitive programme and show that astrocyte-targeted overexpression of mitofusin 2 (*Mfn2*) in the NAc is sufficient to reduce anxiety-like behaviour. Together, these findings identify a testosterone-sensitive neurovascular programme in the NAc linking low testosterone to anxiety vulnerability and implicate astrocytic Mfn2 as a functional component.

## Results

### Low endogenous testosterone is associated with and contributes to high-anxiety-like behaviour in male rats

To determine whether endogenous testosterone is linked to naturally occurring variation in anxiety-like behaviour, we used an established outbred Wistar rat model in which males are selected at the tails of trait anxiety distribution, i.e., as low-anxiety (LA) and high-anxiety (HA) phenotypes, using a validated battery of exploratory anxiety-related assays, including the elevated plus maze (EPM), open field (OF) and novel object (NO) tests (**Fig. 1a**)^50–53^. HA rats displayed reduced open-arm exploration in the EPM, reduced centre exploration in the open field and reduced exploration of the novel object (**Fig. 1b-d**). Integration of these behavioural measures into a composite anxiety z-score captured the robust behavioural separation between the groups (**Fig. 1e**). We next measured plasma testosterone after behavioural phenotyping (**Fig. 1f**). HA rats showed significantly lower plasma testosterone levels than LA rats, and testosterone levels were inversely correlated with the composite anxiety z-score (**Fig. 1f**). Thus, naturally occurring high-anxiety-like behaviour in male rats is associated with low endogenous testosterone.

**Fig. 1:**
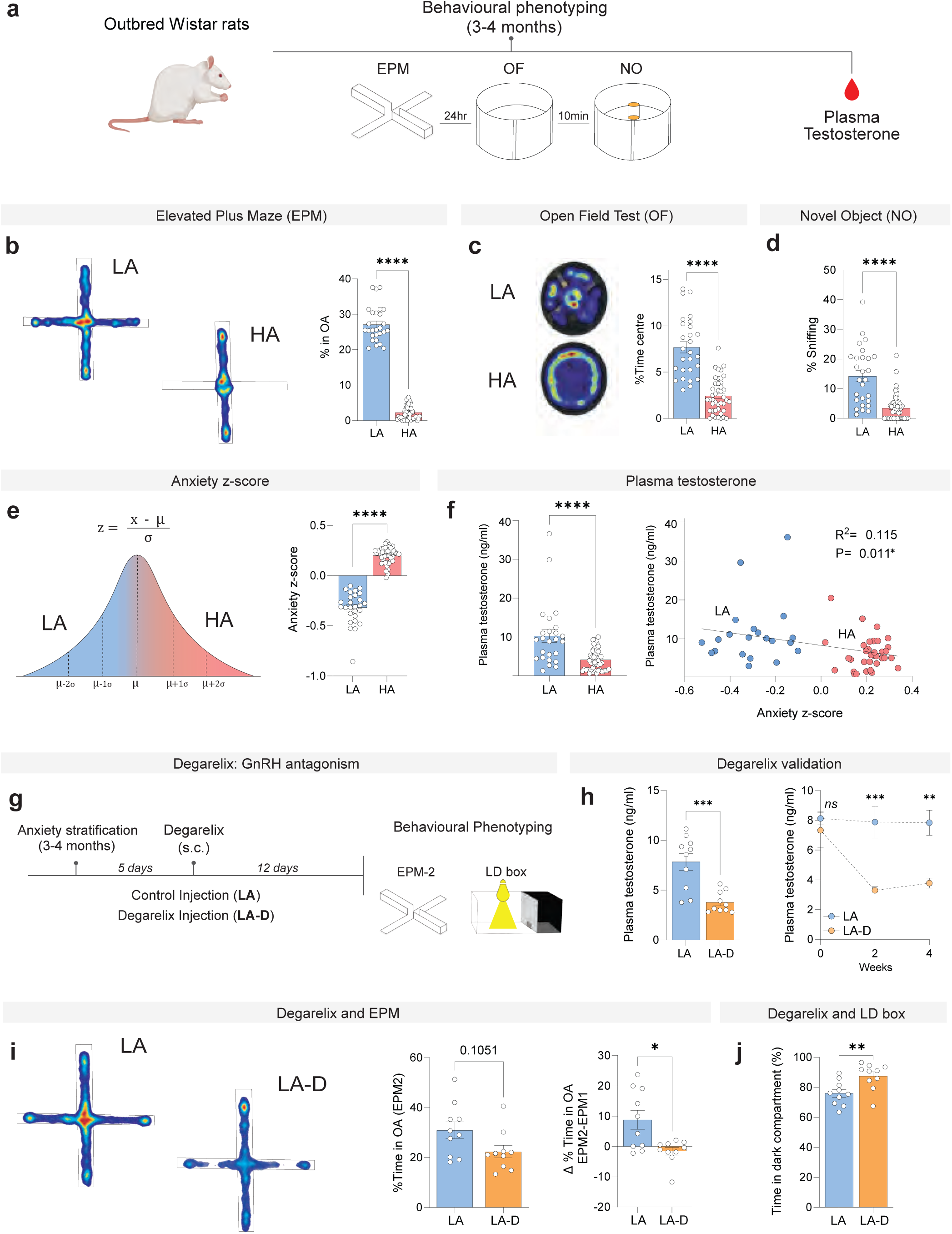
Low endogenous testosterone is associated with and contributes to high anxiety-like behaviour in male rats. **a**, Experimental design for anxiety phenotyping in outbred Wistar rats. *EPM*: elevated plus maze, *OF*: open field test; *NO*: novel object test. **b**, Representative EPM heat maps from low-anxiety (LA) and high-anxiety (HA) rats and percentage of time spent in the open arms (LA, n = 27; HA, n = 43; two-tailed Mann-Whitney U test, *p*<0.0001). **c**, Representative OF heat maps from LA and HA rats and percentage of time spent in the centre (LA, n = 27; HA, n = 43; two-tailed Mann-Whitney U test, *p*<0.0001). **d**, Percentage of time spent sniffing the novel object (LA, n = 27; HA, n = 55; two-tailed Mann-Whitney U test, *p*<0.0001). **e**, Schematic of the composite anxiety z-score and scores in LA and HA rats (LA, n = 27; HA, n = 43; two-tailed Mann-Whitney U test, *p*<0.0001). **f**, Plasma testosterone concentrations in LA and HA rats (LA, n = 25; HA, n = 50; two-tailed Mann-Whitney U test, *p*<0.0001) and Pearson correlation of plasma testosterone concentration with the anxiety z-score (n = 55, r = -0.339, R^2^ = 0.115, two-tailed *p* = 0.011). **g**, Experimental design for pharmacological testosterone suppression in LA rats using the gonadotropin-releasing hormone (GnRH) receptor antagonist degarelix administered subcutaneously (s.c.), followed by behavioural testing in the EPM and light-dark (LD) box. **h**, Plasma testosterone concentrations in vehicle-treated LA rats and degarelix-treated LA rats (LA-D) 4 weeks post-injection (LA, n = 10; LA-D, n = 10; unpaired two-tailed Student’s t-test, *p* = 0.0007) and longitudinal plasma testosterone concentrations over 4-weeks after injection (LA, n = 10; LA-D, n = 10; two-way repeated measures ANOVA; *p*= 0.0013). **i**, Representative EPM heat maps from LA and LA-D rats and percentage of time spent in the open arms during the second EPM exposure (EPM2) (LA, n = 10; LA-D, n = 10; two-tailed Mann-Whitney U test, *p*=0.105). Change (Δ) in the percentage of time spent in the open arms (EPM2-EPM1) (LA, n = 10; LA-D, n = 10; two-tailed Mann-Whitney U test, *p*=0.036). **j**, Percentage of time spent in the dark compartment of the LD box (LA, n = 10; LA-D, n = 10; unpaired two-tailed Student’s t-test, *p*=0.007). Bar and line plots show mean values; error bars represent s.e.m. Dots represent individual rats. *n* denotes the number of rats. ^✱^*p*<0.05, ^✱✱^*p*<0.01, ^✱✱✱^*p*<0.01 and ^✱✱✱✱^*p*<0.0001.

To test whether lowering endogenous testosterone alters anxiety-like behaviour, we pharmacologically suppressed testosterone production in LA rats using the gonadotropin-releasing hormone receptor (GnRH) antagonist degarelix (**Fig. 1g-j**). After anxiety stratification, LA rats received a single subcutaneous injection of degarelix or vehicle and were retested 12 days later in the EPM. They were also tested in the light-dark box (LD), an anxiety-related assay not used during the initial stratification to account for any habituation to exposure. Degarelix markedly reduced plasma testosterone at all post-treatment time points examined, including 4 weeks after treatment (**Fig. 1h**), accompanied by reduced body-weight gain and a lower testis-to-body-weight ratio, consistent with suppression of the hypothalamic-pituitary-gonadal axis (**Extended Data Fig. 1a,b**). Behaviourally, the change in open-arm exploration from the initial phenotyping exposure (EPM1 to EPM2) was significantly lower in degarelix-treated rats than in vehicle-treated controls (**Fig. 1i**). In the LD box, degarelix-treated rats spent significantly more time in the dark compartment (**Fig. 1j**). Together, these findings link naturally low endogenous testosterone to high-anxiety-like behaviour and show that pharmacological suppression of testosterone shifts low-anxiety rats towards higher anxiety-like responding.

### High-anxiety male rats display an NAc-selective increase in BBB permeability and astrocytic endfoot mitochondrial alterations

Having established that HA rats show reduced circulating testosterone levels and that testosterone suppression increases anxiety-like responding, we next asked whether this high-anxiety phenotype was accompanied by alterations at the neurovascular interface (**Fig. 2**). After behavioural stratification, we first mapped BBB permeability across candidate brain regions using Evans Blue injections, an albumin-binding tracer with limited brain parenchymal accumulation when the BBB is intact (**Fig. 2a, b**). HA rats showed significantly higher Evans Blue accumulation in the NAc, whereas no differences were detected in the dorsal striatum (DS) or medial prefrontal cortex (mPFC). Moreover, Evans Blue accumulation was significantly associated with the composite anxiety z-score in the NAc, but not in the other brain regions examined (**Fig. 2b**). Thus, high-anxiety-like behaviour in male rats was associated with a regionally selective increase in BBB permeability in the NAc. Together with prior evidence implicating the NAc in stress-sensitive BBB regulation^47,48,54^ and anxiety-related neurocircuitry^51,52,55,56^, this regionally selective permeability phenotype provided a biologically grounded rationale for examining BBB-associated ultrastructure and astrocytic endfoot organisation in the NAc.

**Fig. 2:**
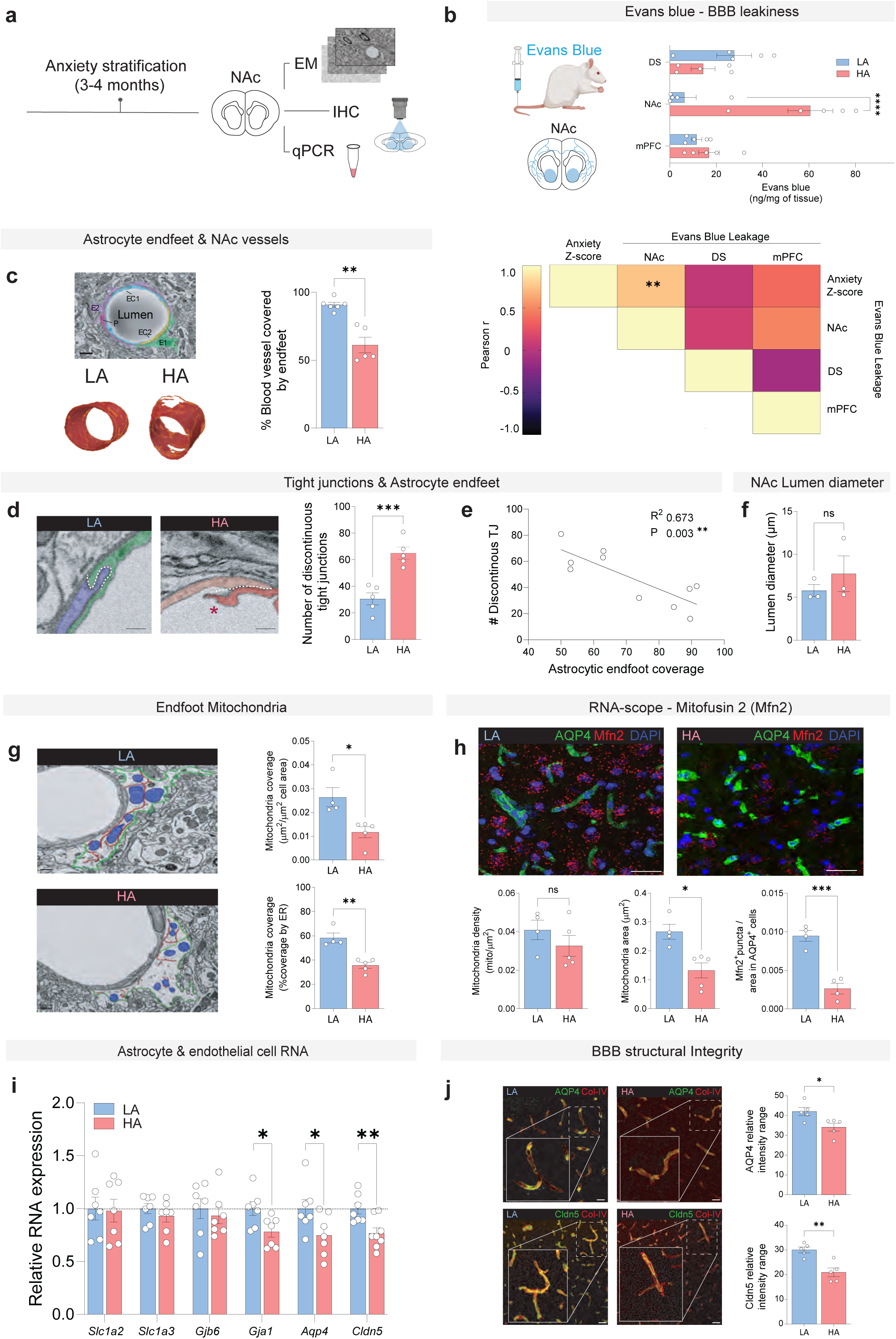
High anxiety male rats display and NAc-selective increase in BBB permeability and altered astrocytic endfoot mitochondrial organisation. **a,** Experimental design for anxiety stratification followed by ex vivo nucleus accumbens (NAc) collection for electron microscopy (EM), immunohistochemistry (IHC) and qPCR analyses. **b,** Schematic of the Evans blue BBB permeability assay and quantification of Evans blue accumulation in the dorsal striatum (DS), NAc and medial prefrontal cortex (mPFC) of LA and HA rats (LA, n = 5; HA, n = 5; two-way ANOVA; anxiety phenotype x brain region interaction, *p<*0.0001; Šídák’s multiple comparisons; mPFC; *p* =0.903, NAc; *p* <0.0001, DS; *p=*0.361). Pearson correlation matrix between anxiety z-score and Evans blue accumulation across the three brain regions; the correlation for NAc was significant (n = 10; r = 0.756, two-tailed *p*=0.0115). **c,** Representative electron microscopy image of an NAc microvessel, with endothelial cells (EC1 and EC2), pericyte (P), and astrocytic endfeet (E1 and E2) colour-coded, and representative 3D reconstructions of astrocytic endfoot coverage of NAc vessels in LA and HA rats. Percentage of the vascular surface covered by astrocytic endfeet (LA, n = 6; HA, n = 5; unpaired two-tailed Student’s t-test, *p*=0.005), Scale bar = 1µm. **d,** Representative EM images of endothelial tight junction morphology in LA and HA rats and quantification of the number of discontinuous tight junctions (LA, n = 5; HA, n = 5; unpaired two-tailed Student’s t-test, *p*=0.0008), Scale bar = 500nm, ✱ on HA signifies discontinuous tight junction. **e**, Pearson correlation between the number of discontinuous tight junctions and astrocytic endfoot coverage (n = 10; r = -0.821, two-tailed *p*=0.0036). **f**, NAc vessel lumen diameter in LA and HA rats (LA, n = 3; HA, n = 3; two-tailed Mann-Whitney U test, *p*=0.400). **g**, Representative EM images of astrocytic endfeet associated mitochondria in the NAc-. Green outlines delineate the astrocytic endfoot boundary, endoplasmic reticulum (ER) is shown in red and mitochondria in blue. Scale bar, 500 nm. Quantification of mitochondrial coverage within astrocytic endfeet and mitochondria-ER coverage (LA, n = 4; HA, n = 4; unpaired two-tailed Student’s t-tests, *p*=0.026 and *p*=0.0015, respectively). **h**, Representative NAc images combining AQP4 immunofluorescence (green) with RNAscope fluorescent *in situ* hybridization (FISH) for Mfn2 mRNA (red) and DAPI nuclear staining (blue). Scale bar, 25 µm. Quantification of astrocytic endfoot mitochondrial density (LA, n = 4; HA, n = 5; Mann-Whitney U test, *p*=0.286), mean mitochondrial area (LA, n = 4; HA, n = 5; Mann-Whitney U test, *p*=0.0159), and Mfn2-positive mRNA puncta per AQP4-positive area (LA, n = 4; HA, n = 4; two-tailed Student’s t-test, *p*=0.0004). **i,** Relative NAc mRNA expression of astrocyte- and BBB-associated genes in LA and HA rats (LA, n 0 7; HA. n = 7; unpaired two-tailed Student’s t-tests): *Slc1a2* (GLT-1) (*p*=0.906); *Slc1a3* (GLAST) (*p*=0.365); *Gjb6* (Cx30) (*p*=0.596); *Gja1* (Cx43) (*p*=0.022); *Aqp4* (*p*=0.047), *Cldn5* (*p*=0.007). **j**, Representative immunofluorescence images of AAP4 and CLDN5 together with Col-IV in the NAc of LA and HA rats. Scale bar, 50 µm. Quantification of AQP4 coverage of Col-IV-positive vessels (LA, n = 5; HA, n = 5; unpaired two-tailed Student’s t-test, *p*=0.025) and CLDN5 (LA, n = 5; HA, n = 5; unpaired two-tailed Student’s t-test, *p*=0.0027). Bar plots show mean values; error bars represent s.e.m. Dots represent individual rats. ns, not significant; ^✱^*p*<0.05, ^✱✱^*p*<0.01, ^✱✱✱^*p*<0.01 and ^✱✱✱✱^*p*<0.0001.

Serial electron microscopy (EM) analyses of NAc microvessels revealed a significant reduction in the percentage of the vascular surface covered by astrocytic endfeet in HA rats (**Fig. 2c**). This was accompanied by a greater number of endothelial tight junction discontinuities (**Fig. 2d**), and the number of discontinuities was inversely correlated with astrocytic endfoot coverage across animals (**Fig. 2e**). These differences were not explained by altered vessel calibre, as lumen diameter did not differ between LA and HA rats (**Fig. 2f**). Moreover, 3D reconstructions did not reveal significant differences in endothelial cell area or volume, pericyte cell area or volume, or endothelial mitochondrial density, surface area or coverage (**Extended Data Fig. 2**). Thus, high-anxiety-like behaviour was associated with reduced astrocytic endfoot coverage and altered tight-junction continuity at NAc vessels, without evidence of broader morphological changes in endothelial cells or pericytes in the same ultrastructural datasets.

Astrocytic endfeet are specialised perivascular domains that contribute to BBB properties, vascular signalling and exchange between blood and brain, and their molecular organisation is essential for neurovascular unit function^40,57^. We therefore examined whether the astrocytic endfoot phenotype was accompanied by changes in mitochondrial organization. HA rats showed reduced mitochondrial coverage within astrocytic endfeet and reduced mitochondria-endoplasmic reticulum (ER) contacts (**Fig. 2g**). Although mitochondrial density was not significantly altered, mean mitochondrial area was reduced in HA rats (**Fig. 2h**), indicating altered mitochondrial morphology and distribution within astrocytic endfeet. Given the regulatory role of mitofusin 2 (Mfn2) in mitochondrial fusion and mitochondria-ER contacts^58,59^, we next combined RNAscope detection of *Mfn2* mRNA with aquaporin 4 (AQP4) immunohistochemistry to quantify *Mfn2*-positive puncta within AQP4-positive (i.e., perivascular) astrocytic domains. HA rats showed fewer *Mfn2*-positive puncta within AQP4-positive domains of the NAc (**Fig. 2h**), consistent with the ultrastructural evidence for altered endfoot-associated mitochondrial organization. In contrast, parallel analyses of endothelial cells did not reveal differences in mitochondrial density, surface area or coverage between LA and HA rats (**Extended Data Fig. 2b**). Pericyte structure was also unchanged (**Extended Data Fig. 2c**). Thus, among the neurovascular compartments examined, the mitochondrial phenotype was detected in astrocytic endfeet but not in endothelial cells and was not accompanied by broader structural alterations in pericytes.

To determine whether these structural and mitochondrial changes were accompanied by molecular alterations in neurovascular markers, we performed qPCR on bilateral NAc punches. HA rats showed reduced expression of *Gja1* (connexin-43, an astrocytic gap-junction protein), *Aqp4* and *Cldn5* (Claudin-5, an endothelial tight-junction protein), whereas *Slc1a2* (GLT-1), *Slc1a3* (GLAST) and *Gjb6* (Cx30) were unchanged (**Fig. 2i**). To determine whether the changes in *Aqp4* and *Cldn5* transcripts were reflected at the protein level, we co-stained NAc sections for AQP4 or CLDN5 together with collagen IV to delineate vascular profiles (**Fig. 2j**). Consistent with the qPCR data, HA rats showed reduced AQP4 coverage of collagen IV-positive vessels and lower vessel-associated CLDN5 signal (**Fig. 2j**). Together with the ultrastructural findings, these results indicate that HA rats display altered astrocytic endfoot organization, reduced endfoot-associated mitochondrial features and lower expression of BBB-associated markers in the NAc.

Finally, we examined whether the relationship between anxiety-like behaviour and BBB permeability observed in males was also detectable in female Wistar rats. Although females could be separated on the basis of EPM open-arm exploration, this separation did not extend consistently to the OF or NO tests and did not yield the robust composite behavioural segregation observed in males (**Extended Data Fig. 3)**. Evans Blue accumulation did not differ between female groups in the regions examined and did not correlate with anxiety-related behavioural measures (**Extended Data Fig. 4**). Complementary analyses stratifying females according to high versus low Evans Blue accumulation likewise did not identify corresponding behavioural differences (**Extended Data Fig. 4**). Thus, with the behavioural and permeability readouts used here, the relationship between anxiety-like behaviour and BBB permeability that was evident in the male NAc was not detected in females. This absence of a detectable female NAc phenotype is consistent with the broader view that stress-related BBB alterations are organised in a region- and sex-dependent manner across affective and mood-related circuits^49,60^. Nevertheless, the absence of a detectable anxiety-related BBB permeability phenotype in females should not be interpreted as evidence against sex specific neurovascular involvement in anxiety, particularly because females did not show the robust behavioural segregation observed in males. We therefore focused subsequent mechanistic experiments on males, where low circulating testosterone levels, high anxiety like behaviour and NAc BBB alterations converged.

Together, these findings identify a NAc neurovascular phenotype in high-anxiety male rats, characterised by increased BBB permeability, reduced astrocytic endfoot coverage, a greater number of tight-junction discontinuities, altered endfoot-associated mitochondrial organisation, reduced *Mfn2* signal in AQP4-positive domains and lower expression of BBB-associated markers. These findings provided the basis for testing whether increasing circulating testosterone in HA rats could shift the NAc neurovascular phenotype towards that observed in low-anxiety rats while reducing anxiety-like behaviour.

### Testosterone supplementation reconfigures mitochondrial and BBB-related transcriptional programmes in NAc astrocytes

Having found that HA rats show reduced circulating testosterone levels together with NAc astrocytic neurovascular alterations, we next tested whether increasing circulating testosterone in HA rats could shift their NAc astrocytic phenotype towards that observed in LA rats (**Fig. 3a**). HA rats were implanted subcutaneously with a slow-release testosterone pellet (HA-T) or control pellet (HA-CTR) and underwent behavioural testing 12 days later, followed by NAc tissue collection for single-nucleus RNA sequencing (snRNA-seq) and immunohistochemical analyses (**Fig. 3a**). Testosterone pellet implantation increased plasma testosterone levels in HA-T rats, validating the supplementation protocol (**Fig. 3b**). We first analysed the snRNA-seq data to define the astrocytic transcriptional response to testosterone supplementation.

**Fig. 3:**
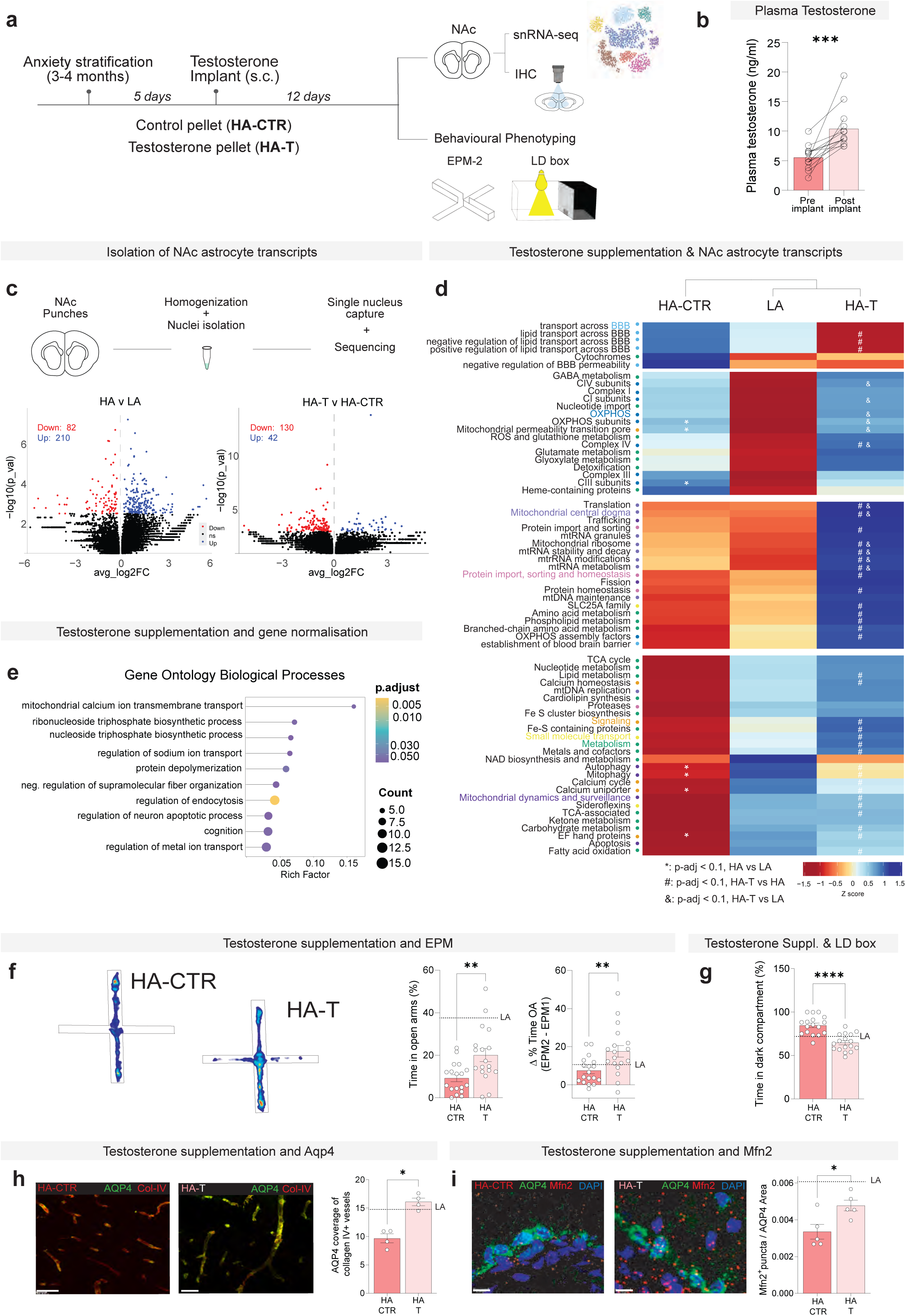
Testosterone supplementation reconfigures NAc astrocytic mitochondrial and BBB-related transcriptional programs and attenuates anxiety-like behaviour in HA rats. **a**, Experimental design for testosterone supplementation in HA rats. HA rats received a subcutaneous (s.c.) control pellet (HA-CTR) or testosterone pellet (HA-T), followed by behavioural testing and NAc collection for snRNA-seq and immunohistochemical analyses. **b**, Plasma testosterone concentrations before and after testosterone pellet implantation in HA-T rats (n = 13; two-tailed Wilcoxon matched-pairs signed-rank test, *p*=0.0002). **c,** Schematic of NAc tissue processing for snRNA-seq (top) and volcano plots of differential gene expression in NAc astrocytes (bottom) for HA-CTR (n = 5 rats) versus LA (n = 5 rats) (left) and HA-T (n = 5 rats) versus HA-CTR (n = 5 rats) (right). Differential expression was assessed using MAST; genes with *p* < 0.01 are shown in red (downregulated) or blue (upregulated), and non-significant genes in black. **d,** Heatmap showing scaled mean mitochondrial and BBB-related pathway scores in NAc astrocytes from HA-CTR, LA and HA-T rats. Columns are hierarchically clustered and rows are grouped by *k*-means clustering (k = 4). Pathway names are colour-coded by encompassing category: BBB, cyan; OXPHOS, blue; mitochondrial central dogma, light purple; protein import, sorting and homeostasis, pink; signalling, orange; small-molecule transport, yellow; metabolism, green; mitochondrial dynamics and surveillance, dark purple. ^✱^ denote adjusted value (p-adj with Benjamini-hochberg corrections) p< 0.1 for HA versus LA, # p-adj < 0.1 for HA-T vs HA, &-p-adj < 0.1, HA-T vs LA. **e,** Gene ontology (GO) enrichment analysis Biological Process enrichment analysis of HA-associated astrocytic genes shifted towards the in LA transcriptional state following testosterone supplementation (i.e., in HA-T). These genes were defined as differing between HA-CRT and LA astrocytes (*p*<0.01). Dot size represents the number of genes and colour represents the adjusted *p* value. Rich factor denotes the proportion of genes in each GO term represented in the analysed gene set with. **f**, Representative EPM heat maps from HA-CTR and HA-T rats and percentage of time in the open arms during the second EPM exposure (EPM2) (HA-CTR, n = 18; HA-T, n = 18; unpaired two-tailed Student’s *t-test*, *p*=0.0039). Change (Δ) in the percentage of time spent in the open arbs between EPM1 and EPM2 (HA-CTR, n = 18; HA-T, n = 18; unpaired two-tailed Student’s *t-test*, *p*=0.0061). **g**, Percentage of time spent in the dark compartment of the LD box (HA-CTR, n = 17; HA-T, n = 17; unpaired two-tailed Student’s *t-test*, *p*< 0.0001). **h**, Representative immunofluorescence images of AQP4 and Col-IV in the NAc of HA-CTR and HA-T rats. Quantification of AQP4 coverage of Col-IV-positive vessels (HA-CTR, n = 4; HA-T, n = 4; two-tailed Mann-Whitney U test, *p*=0.0286). Scale bar, 50 µm. **i,** Representative NAc images combining AQP4 immunofluorescence (green) with RNAscope fluorescent *in situ* hybridization (FISH) for *Mfn2* mRNA (red) and DAPI nuclear staining (blue) in HA-CTR and HA-T rats. Quantification of *Mfn2*-positive mRNA puncta in AQP4-positive area (HA-CTR, n = 5; HA-T, n = 5; unpaired two-tailed Student’s t-test, *p*=0.0215). Scale bar, 10 µm. Dashed horizontal lines in f-I indicate the corresponding mean values in LA rats for reference. For b and f-I, dots represent individual rats. Bar plots show mean values, error bars represent s.e.m. n denotes the number of rats unless otherwise stated. ns, not significant; ^✱^*p* < 0.05, ^✱✱^*p*< 0.01, ^✱✱✱^*p*<0.01 and ^✱✱✱✱^*p*< 0.0001.

We isolated nuclei from the NAc of HA-CTR, LA and HA-T animals and carried out snRNA-seq. Unsupervised clustering recovered the major NAc cell types previously reported^53^ (**Extended Data Fig. 5a**). Focusing on astrocytes, we identified 292 differentially expressed genes between HA-CTR and LA, and 172 between HA-CTR and HA-T rats (**Fig. 3c**). To determine whether the HA phenotype and testosterone supplementation were associated with changes in mitochondrial and BBB-related pathways in astrocytes, we applied a pathway-scoring framework based on an in silico mitotyping approach^53,61^. Expression values of nuclear-encoded mitochondrial genes were aggregated into 149 mitopathway scores based on MitoCarta 3.0^62^, and genes associated with BBB functions were aggregated into 14 pathway scores based on Gene Ontology annotations. K-means clustering of the pathway scores across the groups highlighted a shift in HA-T astrocytes towards an LA-like pathway profile (**Fig. 3d**). Compared with LA astrocytes, HA-CTR astrocytes showed higher scores for OXPHOS-related pathways, including OXPHOS, complex III and complex IV subunits, and the mitochondrial permeability transition pore, together with lower scores for mitophagy, autophagy, and calcium-related pathways, including the calcium uniporter and EF-hand proteins (**Fig. 3d**). Relative to HA-CTR astrocytes, testosterone supplementation reduced scores for BBB-related lipid transport pathways and increased scores for pathways related to mitochondrial gene expression, mitochondrial dynamics and quality control, intermediary metabolism, and signalling and calcium regulation (**Fig. 3d**). Among pathways that differed both between HA-CTR and LA astrocytes and between HA-CTR and HA-T astrocytes, testosterone supplementation shifted a subset related to mitochondrial dynamics and calcium regulation towards the LA profile, whereas OXPHOS-related differences did not show a comparable shift. HA-T astrocytes also exhibited changes in other metabolic pathways that were not altered between HA-CTR and LA astrocytes, indicating that the transcriptional effects of testosterone were not limited to reversal of the HA-associated profile. Together, this pathway-level analysis indicates that testosterone supplementation reconfigures NAc astrocytic transcriptional programmes linked to mitochondrial dynamics, calcium handling, metabolism and BBB-associated functions.

We next quantified the proportion of HA-associated astrocytic genes that were no longer significantly different from LA astrocytes after testosterone treatment. Specifically, this criterion comprised genes that differed between HA-CTR and LA astrocytes (*p* < 0.01) but not between HA-T and LA astrocytes (i.e., shifted towards the LA transcriptional state). In line with the findings from the mitotyping analysis, 70.2% of the HA-associated genes met this criterion in the HA-T astrocytes (**Extended Data Fig. 5b**). Gene Ontology analysis of this subset highlighted mitochondrial calcium ion transmembrane transport, ribonucleoside and nucleoside triphosphate biosynthetic processes, regulation of sodium and metal ion transport, regulation of endocytosis, and cytoskeletal organisation-related terms (**Fig. 3e**). Among the genes contributing to these categories were mitochondrial calcium-handling genes, including *Slc8b1*, *Vdac1*, *Micu1* and *Micu2*, and vascular- and trafficking-related genes such as *Vegfa* and *Rab4b* (**Extended Data Fig. 5c-e).** Together, these results identify a testosterone-sensitive astrocytic transcriptional programme in the NAc, enriched for mitochondrial calcium signalling, mitochondrial metabolism and mitochondrial dynamics.

### Testosterone supplementation attenuates anxiety-like behaviour and shifts NAc astrocytic and BBB features towards the LA profile

Having found that testosterone supplementation shifted a subset of HA-associated NAc astrocytic transcriptional programmes towards the LA profile, we next asked whether supplementation also reduced anxiety-like behaviour in HA rats. In the EPM, HA-T rats spent significantly more time in the open arms than HA-CTR rats and showed a larger increase in open-arm exploration relative to their initial phenotyping exposure (**Fig. 3f**). In the LD box, HA-T rats spent significantly less time in the dark compartment than HA-CTR rats (**Fig. 3g**). Comparisons with LA animals indicated that testosterone supplementation shifted the HA behavioural phenotype towards that of LA rats (**Extended Data Fig. 6a,b**). Thus, testosterone supplementation attenuated anxiety-like behaviour in HA rats.

We next examined whether testosterone supplementation also modified the astrocytic endfoot and mitochondrial features altered in HA rats. Testosterone significantly increased AQP4 coverage of collagen IV-positive vessels in the NAc (**Fig. 3h**). In parallel, HA-T rats showed more *Mfn2*-positive puncta within AQP4-positive domains (**Fig. 3i**), indicating that testosterone shifted the astrocytic endfoot-associated Mfn2 phenotype towards the LA profile. Direct comparisons with LA animals showed that AQP4 coverage in HA-T animals was shifted towards LA levels and *Mfn2* also showed a partial shift towards the LA profile (**Extended Data Fig. 6c**). CLDN5 levels showed a similar direction of change, but the difference between HA-CTR and HA-T groups did not reach significance (**Extended Data Fig. 6d**). Given that the HA neurovascular phenotype was also characterised by increased Evans Blue accumulation in the NAc, we next asked whether testosterone supplementation also affected this BBB permeability readout. HA-T rats showed reduced Evans Blue accumulation in the NAc relative to HA-CTR rats, consistent with reduced NAc BBB permeability (**Extended Data Fig. 6e**). Together, these data indicate that testosterone supplementation shifts key astrocytic endfoot, mitochondrial and BBB permeability features of the HA neurovascular phenotype towards the LA profile.

To complement these in vivo findings with an independent barrier assay, we tested whether testosterone directly modulates endothelial barrier properties in an astrocyte-dependent manner (**Extended Data Fig. 7a**). We measured transendothelial electrical resistance (TEER) in human brain microvascular endothelial cells cultured either alone or in co-culture with primary human astrocytes. Testosterone did not significantly alter TEER in endothelial monocultures at any of the concentrations tested (0.1, 5, and 100 nM) (**Extended Data Fig. 7b**). In contrast, testosterone increased TEER at 0.1 and 5 nM in endothelial-astrocyte co-cultures (**Extended Data Fig. 7c**). Furthermore, flutamide, an AR antagonist, decreased TEER in endothelial-astrocyte co-cultures, whereas it had no significant effect in endothelial monocultures (**Extended Data Fig. 7d**). These in vitro data support an astrocyte-dependent effect of androgen signalling on endothelial barrier resistance.

In sum, testosterone supplementation attenuated anxiety-like behaviour in HA rats and shifted a subset of NAc astrocytic transcriptional programmes, perivascular AQP4 coverage, endfoot-associated Mfn2 and BBB permeability towards the LA profile. The co-culture findings further support an astrocyte-dependent effect of androgen signalling on endothelial barrier resistance. We next asked whether similar testosterone-sensitive NAc astrocytic features were evident in an independent condition of endogenous low testosterone.

### Testosterone supplementation reduces anxiety-like behaviour and increases perivascular AQP4 and Mfn2 in the NAc of older male rats

Because circulating testosterone levels decline with age in males^13,14^, including in male rats^63^, ageing provides an independent condition of endogenous low testosterone in which to test the broader relevance of the testosterone-sensitive NAc astrocytic phenotype. We therefore examined older male rats to determine whether testosterone supplementation would reduce anxiety-like behaviour and engage the same NAc astrocytic features that were testosterone-sensitive in HA rats (**Fig. 4a**). Eleven-month-old male rats showed significantly lower plasma testosterone levels than 3-month-old males (**Fig. 4b**). We then implanted older rats with either a slow-release testosterone pellet (Old-T) or a control pellet (Old-CTR) and confirmed that testosterone supplementation elevated plasma testosterone levels for at least 30 days after implantation, reaching levels similar to those in 3-month-old males (**Extended Data Fig. 8**).

**Fig. 4:**
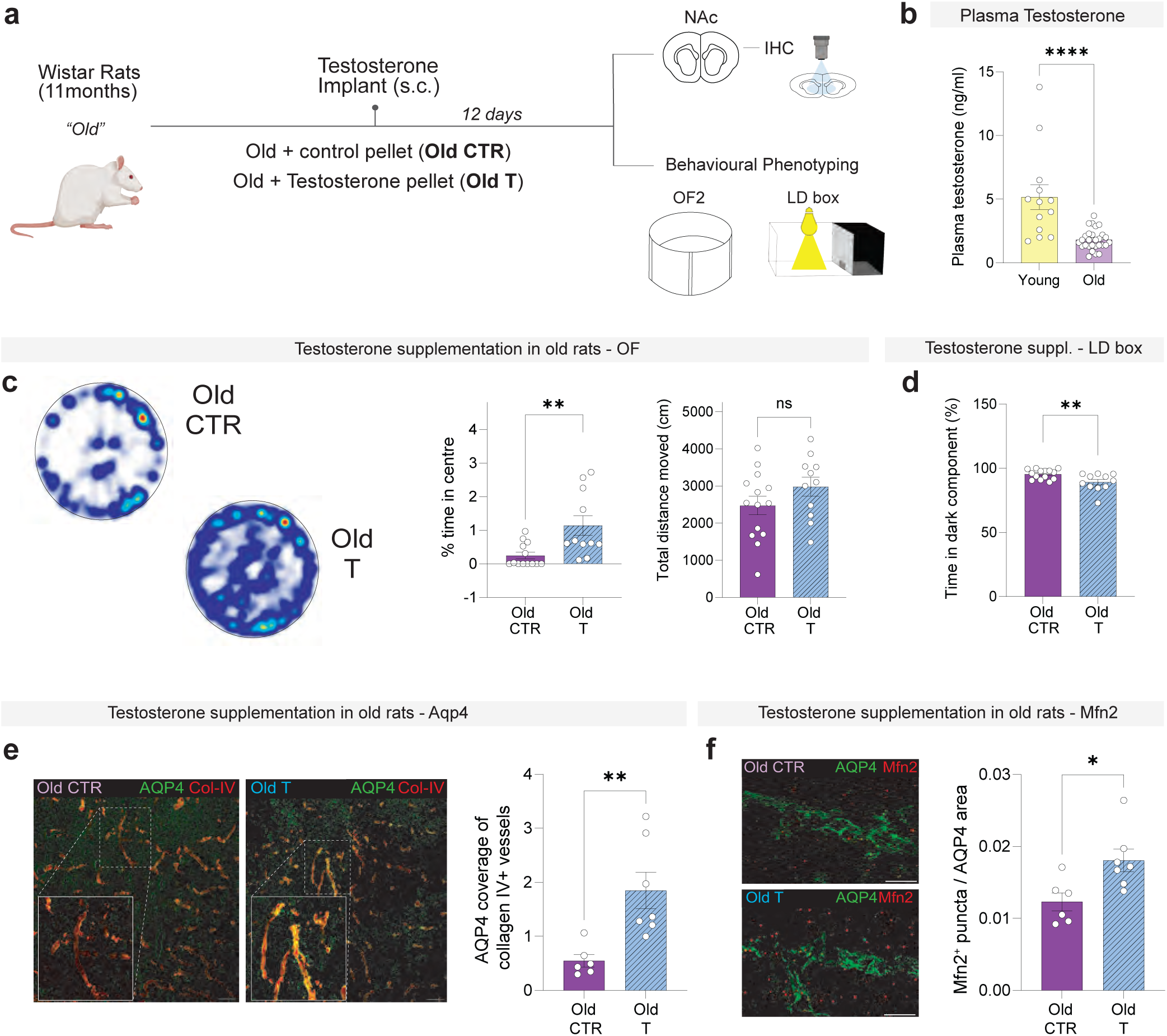
Testosterone supplementation reduces anxiety-like behaviour and increases NAc perivascular AQP4 and Mfn2 in older male rats. **a**, Experimental design for testosterone supplementation in 11-month-old male Wistar rats. Rats received a subcutaneous (s.c.) control pellet (Old-CTR) or testosterone pellet (Old-T), followed 12 days later by behavioural testing and NAc collection for immunohistochemical analyses. **b**, Plasma testosterone concentration in 3-month-old (young) and 11-month-old (older) male rats (young, n = 13; older, n = 28; unpaired two-tailed Student’s t-test, *p*< 0.0001). **c,** Representative OF heat maps from Old-CTR) and Old-T rats. Percentage of time spent in the centre (Old-CTR, n = 13; Old-T, n = 11; unpaired two-tailed Student’s t-test, *p*=0.004) and total distance moved (Old-CTR, n = 14; Old-T, n = 11; unpaired two-tailed Student’s t-test, *p*=0.172). **d,** Percentage of time spent in the dark compartment of the LD box (Old-CTR, n = 14; Old-T, n = 11; unpaired two-tailed Student’s t-test, *p*= 0.0089). **e,** Representative immunofluorescence images of AQP4 and Col-IV in the NAc of Old-CTR and Old-T rats. Quantification of AQP4 coverage of C-IV-positive vessels (Old-CTR, n = 6; Old-T, n = 7; two-tailed Mann-Whitney U test, *p*=0.0072). Scale bar, 50 µm. **f**, Representative NAc images combining AQP4 immunofluorescence (green) with RNAscope fluorescent *in situ* hybridization (FISH) for *Mfn2* mRNA (red) and DAPI nuclear staining (blue) in Old-CTR and Old-T rats. Quantification of *Mfn2*-positive mRNA puncta in AQP4-positive area (Old-CTR, n = 6; Old-T, n = 7; unpaired two-tailed Student’s t-test, *p*=0.0157). Scale bar, 50 µm. Bar plots show mean values; error bars represent s.e.m. Dots represent individual rats. n denotes number of rats. ns, not significant; ^✱^*p* < 0.05, ^✱✱^*p*< 0.01, ^✱✱✱^*p*<0.01 and ^✱✱✱✱^*p*< 0.0001.

Twelve days after implantation, Old-CTR and Old-T rats were tested in the OF and LD box; their large body size precluded reliable use of the EPM (**Fig. 4c,d**). Compared with Old-CTR rats, Old-T rats spent significantly more time in the centre of the OF, with no difference in total distance travelled, indicating that the behavioural effect was not explained by altered locomotion (**Fig. 4c**). They also spent significantly less time in the dark compartment of the LD box (**Fig. 4d**). Thus, testosterone supplementation reduced anxiety-like behaviour in older male rats with low circulating testosterone levels.

We next asked whether this behavioural effect was accompanied by changes in the same NAc astrocytic features modified by testosterone in HA rats. Testosterone-treated older rats showed increased AQP4 coverage of collagen IV-positive vessels in the NAc (**Fig. 4e**), together with more *Mfn2*-positive puncta within AQP4-positive domains (**Fig. 4f**). Thus, testosterone supplementation elicited convergent behavioural and NAc astrocytic responses across two distinct conditions of endogenous low testosterone. We next tested whether the behavioural and neurovascular responses to testosterone supplementation in HA rats involve AR signalling within the NAc.

### NAc androgen receptor knockdown attenuates testosterone-dependent behavioural and neurovascular responses in HA rats

To reduce AR signalling within the NAc, we used an adeno-associated viral vector expressing an AR-targeting short hairpin RNA (shRNA) under the U6 promoter, together with GFP. HA rats received bilateral NAc injections of AAV2-GFP-U6-h-AR-shRNA (HA-AR-KD) or a control AAV2-CMV-eGFP virus (HA-GFP), followed by testosterone pellet implantation 20 days later and behavioural testing in the EPM and LD box 12 days after implantation (**Fig. 5a**).

**Fig. 5:**
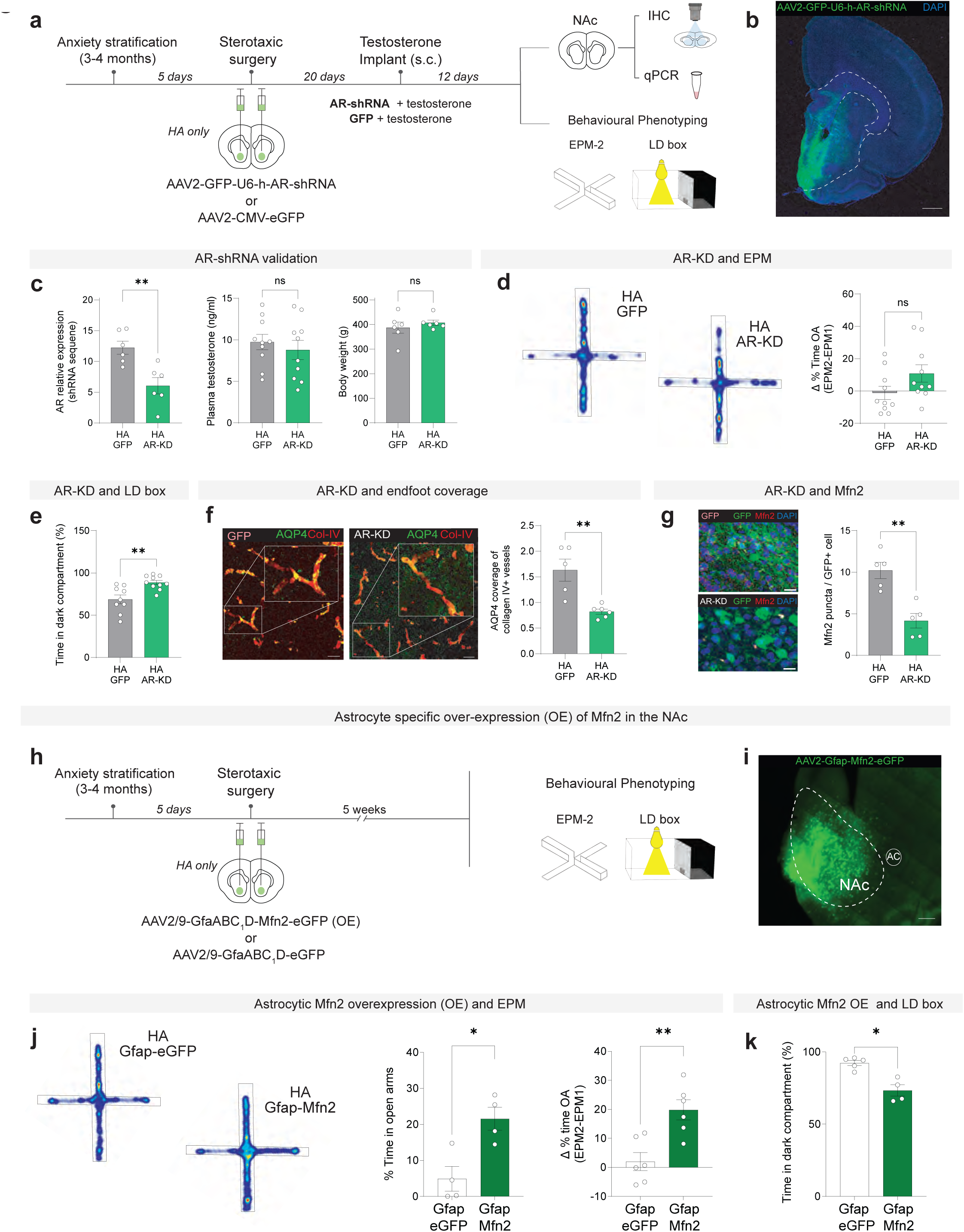
NAc androgen receptor signalling contributes to testosterone-dependent behavioural and astrocytic responses, while astrocytic Mfn2 overexpression reduces anxiety-like behaviour. **a**, Experimental design for NAc androgen receptor (AR) knockdown (KD) in HA rats. Rats received bilateral NAc injections of AAV2-GFP-U6-h-AR-shRNA (HA-AR-KD) or control (AAV2-CMV-eGFP (HA-GFP), followed 20 days later by subcutaneous (s.c.) testosterone pellet implantation and from 12 days later behavioural testing. **b**, Representative image showing viral GFP expression following of AAV2-GFP-U6-h-AR-shRNA delivery to the NAc. Scale bar, 100 µm. **c**, Validation of viral-mediated AR KD by relative AR expression in NAc punches from HA-GFP and HA-AR-KD rats (HA-GFP, n = 6; HA-AR-KD, n = 6; unpaired two-tailed Student’s t-test, *p*=0.005). Plasma testosterone levels following testosterone supplementation (HA-GFP, n = 10; HA-AR-KD, n = 10; unpaired two-tailed Student’s t-test, *p*=0.527) and body weight (HA-GFP, n = 6; HA-AR-KD, n = 6; unpaired two-tailed Student’s t-test, *p*=0.444). **d**, Representative EPM heat maps from HA-GFP and HA AR-KD rats and change (Δ) in the percentage of time spent in the open arms (EPM2-EPM1) (HA-GFP, n = 10; HA-AR-KD, n = 10; unpaired two-tailed Student’s t-test, *p*=0.091). **e**, Percentage of time spent in the dark compartment of the LD box (HA-GFP, n = 9; HA-AR-KD, n = 10; unpaired two-tailed Student’s t-test, *p*=0.003). **f**, Representative immunofluorescence images of AQP4 and Col-IV in the NAc of HA-GFP and HA-AR-KD rats. Quantification of AQP4 coverage of C-IV-positive vessels (HA-GFP, n = 5; HA-AR-KD, n = 6; unpaired two-tailed Student’s t-test, *p*=0.003. Scale bar, 50µm. **g**, Representative NAc images showing GFP-positve virally transduced cells, RNAscope fluorescent *in situ* hybridization (FISH) for *Mfn2* mRNA and DAPI nuclear staining in HA-GFP and HA AR-KD rats. Quantification of *Mfn2*-positive mRNA puncta per GFP-positive cell (HA-GFP, n = 5; HA-AR-KD, n = 6; unpaired two-tailed Student’s t-test, *p*=0.002). Scale bar, 10 µm. **h,** Experimental design for astrocyte-targeted *Mfn2* overexpression in the NAc of HA rats. Rats received bilateral NAc injections of AAV2/9-GfaABC_1_D-Mfn2-eGFP (Gfap-Mfn2) or the matched AAV2/9-Gfap-Mfn2-eGFP control (Gfap-eGFP), followed by behavioural testing 5 weeks later. **i,** Representative image showing AAV2/9-GfaABC_1_D-Mfn2-eGFP expression in the NAc of HA rats. Scale bar, 100µm. **j,** Representative EPM heat maps from HA rats expressing Gfap-eGFP or Gfap-Mfn2 and percentage of time spent in the open arms during EPM2 (Gfap-eGFP, n = 4; Gfap-Mfn, n = 4; unpaired two-tailed Student’s t-test, *p*=0.012). Change (Δ) in time the percentage of time spent in the open arms between EPM1 and EPM2 (Gfap-eGFP, n = 6; Gfap-Mfn, n = 6; unpaired two-tailed Student’s t-test, *p*=0.003). **k,** Percentage of time spent in the dark component of the LD box (Gfap-eGFP, n = 5; Gfap-Mfn, n = 4; unpaired two-tailed Student’s t-test, *p*=0.012 Bar plots show mean values; error bars represent s.e.m. Dots represent individual rats. n denotes number of rats. ns, not significant; ^✱^*p* < 0.05, ^✱✱^*p*< 0.01, ^✱✱✱^*p*<0.01 and ^✱✱✱✱^*p*< 0.0001.

We first validated the manipulation in NAc tissue (**Fig. 5b**) and found that AR levels were reduced by approximately 50% in HA-AR-KD rats relative to HA-GFP controls (**Fig. 5c**). NAc AR-KD did not alter plasma testosterone levels or body weight in testosterone-supplemented HA rats (**Fig. 5c**), indicating that subsequent behavioural and cellular differences were not attributable to differences in circulating testosterone or body weight.

Behaviourally, the change in open arm exploration from the initial phenotyping to the post-supplementation EPM did not differ significantly between groups (**Fig. 5d**). By contrast, HA-AR-KD rats spent significantly more time in the dark compartment of the LD box than HA-GFP controls (**Fig. 5e**). Thus, NAc AR signalling contributes to the testosterone-dependent reduction in anxiety-like responding, although the behavioural effect of AR knockdown was detected in the LD box but not in the EPM.

We next asked whether NAc AR-KD attenuated the neurovascular features modified by testosterone supplementation. Compared with HA-GFP controls, HA-AR-KD rats showed reduced AQP4 coverage of collagen IV-positive vessels and lower vessel-associated CLDN5 signal in the NAc (**Fig. 5f; Extended Data Fig. 9a**). HA-AR-KD rats also showed fewer *Mfn2*-positive puncta within AQP4-positive domains (**Fig. 5g**). Together, these findings show that partial reduction of NAc AR levels attenuates testosterone-dependent behavioural and astrocytic responses in HA rats. Because *Mfn2* emerged as a recurrent testosterone-sensitive astrocytic feature and was reduced by NAc AR knockdown, we next tested whether increasing Mfn2 selectively in NAc astrocytes was sufficient to reduce anxiety-like behaviour.

### Astrocyte-targeted Mfn2 overexpression in the NAc is sufficient to attenuate anxiety-like behaviour in HA rats

The preceding experiments identified *Mfn2* as a recurrent testosterone-sensitive feature of the HA NAc astrocytic-neurovascular phenotype. Specifically, *Mfn2* signal in AQP4-positive domains was reduced in HA rats, increased by testosterone supplementation in young HA rats and older low-testosterone males, and the testosterone-associated increase was attenuated by NAc AR knockdown. We therefore asked whether increasing *Mfn2* levels in the NAc using an astrocyte-targeted vector was sufficient to reduce anxiety-like behaviour in HA rats.

To test this hypothesis, we injected HA rats bilaterally into the NAc with an AAV2/9 vector expressing *Mfn2* under the astrocyte-selective GfaABC_1_D promoter (AAV2/9-GfaABC_1_D-Mfn2-eGFP; Gfap-Mfn2) or with a matched control vector (AAV2/9-GfaABC_1_D-eGFP; Gfap-eGFP) (**Fig. 5h**). Viral expression was localised to the NAc and showed the expected astrocyte-targeted expression pattern (**Fig. 5i**). Five weeks after stereotaxic surgery, rats were retested for anxiety-like behaviour in the EPM and LD box (**Fig. 5h, k**). Compared with Gfap-eGFP-injected control rats, Gfap-Mfn2 rats spent significantly more time in the open arms of the EPM and showed a greater increase in open-arm exploration relative to their initial phenotyping exposure (**Fig. 5j**). In the LD box, astrocytic Mfn2 overexpression also reduced the time spent in the dark compartment (**Fig. 5k**).

Thus, increasing *Mfn2* in NAc astrocytes was sufficient to attenuate the high-anxiety behavioural phenotype across two anxiety-related assays. These findings functionally link a recurrent testosterone-sensitive astrocytic mitochondrial feature in the NAc to anxiety-like behaviour.

## Discussion

Here, we identify low endogenous testosterone as a causal endocrine contributor to persistent high anxiety and reveal an NAc neurovascular programme involving mitochondrial remodelling in perivascular astrocytes. By anchoring the study in naturally occurring differences in anxiety-like behaviour in an outbred rat population, we link systemic androgen tone to NAc BBB homeostasis. Convergent behavioural and astrocytic responses in older low-testosterone males support the broader relevance of these NAc astrocytic features across distinct conditions of endogenous low testosterone. Our findings extend circuit-centred models of anxiety by identifying testosterone-sensitive mitochondrial remodelling in perivascular astrocytes as a functional component through which peripheral endocrine state can shape persistent anxiety vulnerability.

A major advance of this study is to place BBB dysfunction in the NAc within the biological architecture of trait anxiety, rather than viewing it solely as a downstream consequence of chronic stress exposure. In outbred male rats with naturally occurring high anxiety, BBB permeability was increased selectively in the NAc, but not in the dorsal striatum or mPFC; NAc permeability was correlated with individual anxiety scores and was accompanied by alterations in tight-junction continuity and perivascular astrocytic organisation. Previous work has shown that chronic stress produces region- and sex-specific BBB alterations linked to stress susceptibility, with prominent involvement of the NAc in males^47–49,54,60,64^. Our findings establish that NAc BBB dysfunction is also present as part of a naturally occurring high-anxiety phenotype, in the absence of an imposed chronic stress paradigm. The basis of this regional selectivity remains unresolved. In females, we did not detect a relationship between BBB permeability and anxiety-related behaviour, although this finding should be interpreted cautiously given the weaker behavioural segregation in this cohort and evidence for sex- and region-dependent neurovascular responses to stress^49,60,65^. In males, our findings further implicate systemic testosterone acting through NAc AR signalling in the regulation of key components of this perivascular phenotype. Because sustained or severe stress can suppress hypothalamic-pituitary-gonadal axis activity and thereby reduce testosterone production^66,67^, our data raise the possibility that reduced androgen signalling may represent one route through which stress converges on the same NAc neurovascular vulnerability.

Within the NAc, a prominent structural feature was reduced astrocytic endfoot coverage rather than broad vascular perturbation. High-anxiety rats showed reduced astrocytic endfoot coverage and perivascular AQP4 coverage, along with more tight-junction discontinuities and lower *Gja1* and *Aqp4* expression. The reduction in *Gja1* is notable given that stress can reduce astrocytic Cx43-mediated coupling and constrain glucose and lactate trafficking through astrocyte networks^68^. Astrocytic endfeet support endothelial barrier properties, water and ion homeostasis, vascular signalling, metabolic exchange and waste clearance^40,57,69,70^, and perivascular astrocytic disorganisation has been described in major depressive disorder and stress models^48,49,60,71^. The contribution of astrocytes to androgen-dependent barrier regulation is further supported by the in vitro BBB assay, in which testosterone increased endothelial barrier resistance only when astrocytes were present, whereas AR antagonism lowered resistance in co-cultures but not in endothelial monocultures. Although the relative contribution of altered barrier permeability itself to behavioural vulnerability remains to be established, the convergence of astrocytic transcriptional and perivascular responses in vivo with the co-culture findings positions astrocytes as central contributors to androgen-sensitive BBB regulation.

A central mechanistic insight from our study is the identification of mitochondrial remodelling within astrocytic endfeet, with *Mfn2* emerging as a functional component of the high-anxiety phenotype. In high-anxiety males, astrocytic endfeet showed reduced mitochondrial coverage and mean mitochondrial area, together with fewer mitochondria–ER contacts and fewer *Mfn2* mRNA puncta in AQP4-positive domains, whereas the endothelial mitochondrial measurements were unchanged. The localisation of these changes to astrocytic endfeet is particularly relevant because these specialised vascular domains couple astrocytic Ca^2+^ signalling to vascular tone^72^ and are enriched in mitochondria-ER contact sites^59^. *Mfn2*-dependent organisation of these contacts regulates mitochondrial Ca^2+^ uptake and perivascular Ca^2+^ signalling and is required for vascular remodelling following cortical injury^59^. This is consistent with the broader role of MFN2, a dynamin-related GTPase involved in mitochondrial dynamics and network organisation, as well as in mitochondria-ER contact organisation^73^. Recent work further links astrocytic mitochondria to neurovascular communication through mitochondrial transfer to endothelial cells and pericytes^74^, with astrocytic *Mfn2* implicated in mitochondrial transfer to endothelial cells and BBB integrity^75^. Consistent with altered mitochondrial Ca^2+^ handling, calcium-related mitopathway scores were reduced in HA astrocytes and a subset shifted towards the LA profile after testosterone supplementation; moreover, HA-associated genes shifted towards the LA transcriptional state were enriched for mitochondrial calcium ion transmembrane transport, including *Slc8b1*, *Vdac1*, *Micu1* and *Micu2*. Together with the reduced mitochondria-ER contacts and *Mfn2* signal, these transcriptional changes point to altered mitochondrial Ca^2+^ handling as a candidate functional consequence of the astrocytic mitochondrial phenotype.

The link between systemic androgen state and this astrocytic mitochondrial phenotype was evident at both transcriptional and cellular levels. Testosterone supplementation reconfigured astrocytic programmes related to mitochondrial dynamics and quality control and increased AQP4-associated *Mfn2* towards the low-anxiety profile, whereas NAc AR knockdown attenuated the increase in *Mfn2*. The regulation of astrocytic mitochondrial programmes by testosterone is consistent with evidence from other tissues that androgen signalling can influence mitochondrial biogenesis and dynamics, including MFN2 expression^76,77^. Evidence in the brain is more limited, but testosterone deprivation has been associated with reduced mitochondrial gene expression and biogenesis in the hippocampus^78^ and impaired mitochondrial respiratory-chain function in the substantia nigra^79^, whereas supplementation was shown to enhance mitochondrial respiration, biogenesis and quality-control programmes^64,80^. Importantly, in our study, astrocyte-targeted *Mfn2* overexpression in the NAc was sufficient to attenuate anxiety-like behaviour, extending evidence for circuit-specific astrocyte signalling in basal ganglia networks^81^ and for functional specialisation of NAc astrocytes in circuit integration and motivated behaviour^82,83^. These findings connect systemic androgen state to *Mfn2*-linked mitochondrial organisation in NAc astrocytes and establish the behavioural relevance of this astrocytic mitochondrial component.

More broadly, mitochondrial regulation has emerged as an important determinant of NAc function and behaviour, including thorough Mfn2-dependent mechanisms in medium spiny neurons^51,52^ and broader alterations in NAc mitochondrial function associated with anxiety, social dominance and motivation^50–53,84,85^. Our findings reveal a complementary cellular and physiological dimension, in which systemic androgen state engages mitochondrial organisation in perivascular astrocytes at the neurovascular interface. This adds to an emerging view in which astrocytic mitochondrial state actively shapes brain homeostasis and behaviour through redox, metabolic and signalling mechanisms^86–88^, and supports a broader framework in which brain mitochondria act as cellular interfaces through which systemic signals can shape circuit function and behaviour^89^.

Importantly, our findings identify and NAc astrocytic mechanism through which low androgen tone can promote anxiety vulnerability, raising the possibility that related mechanisms may be relevant across distinct physiological and clinical contexts. This may help explain the recurrent association of anxiety with lower testosterone in community samples^18^ and clinical hypogonadism^19^, reduced AR signalling in older age^90^, and therapeutic androgen deprivation^20,22^.

In summary, our study places naturally low testosterone within the biology of persistent anxiety vulnerability and identifies the NAc neurovascular interface as a site at which systemic androgen state engages Mfn2-linked astrocytic mitochondrial organisation. These findings broaden models of anxiety beyond neuronal circuits and identify androgen regulation of the astrocytic neurovascular interface as a route through which systemic physiological state can become embedded in brain mechanisms that sustain behavioural vulnerability.

## Methods

### Animals

All procedures were conducted in accordance with the Swiss National Institutional Guidelines on Animal Experimentation and approved by the Swiss Cantonal Veterinary Office Committee for Animal Experimentation. Adult outbred male Wistar rats (Charles River, L’Arbresle, France) were approximately 250g (3-4 months) upon arrival at the facility and were housed in pairs in 2 in polypropylene cages (57x35x20 cm) at 23°C on a 12-hour light-dark cycle (lights-on at 07h00-19h00). All animals had ad libitum access to standard chow and water. Animals were allowed to habituate to the vivarium for one week and were then handled for 2 min/day during 3 days prior to the start of all experiments. Adult female Wistar rats (Charles River, L’Arbresle, France) were age matched to the male rats and maintained under identical housing conditions but in separate rooms. For ageing experiments, rats classified as intermediate-anxious (see *z-score for anxiety*) were maintained under standard housing conditions and allowed to age until 11 months of age; these animals constituted the aged cohort used in the study. All behavioural manipulations were performed during the light phase.

### Behavioural testing

To classify animals into low and high-anxiety phenotypes, we leveraged a validated model^50,51,53^ natural inter-individual variability in anxiety-like behaviours observed within the population using a battery of validated behavioural assays. Anxiety-related behaviour was assessed through spontaneous exploration in the elevated plus maze (EPM), open field test (OF), and novel object tests (NO), followed by the calculation of a composite anxiety z-score. Animals were initially categorized based on their performance in the elevated plus maze, specifically the percentage of time spent in the open arms and classified as high-anxiety (HA; <5% open-arm time) or low-anxiety (LA; >20% open-arm time) as previously described^50,51^. Animals were then further classified based on their exploration of the central zone of the open field and their percentage time sniffing a novel object placed at the center of the arena. Integration of these behavioural parameters into a composite z-score resulted in a robust segregation of animals into HA and LA groups. This score integrates measures of behavioral dimensions over time and tests, yielding a combined measure. A z-score was calculated for each individual animal by computing how many standard deviations σ a given observation X (% time in the open arms for EPM, latency to sniffing, and % time in center for NO) is from the group mean (μ), according to the formula z = (X-μ)/σ. These scores were averaged within each test such that each test had the same weight, and then averaged across different tests (EPM, NO) to obtain a composite z score of anxiety for each animal.

#### Elevated plus maze (EPM)

An elevated platform raised 55 cm above the floor, with two opposite open arms and two opposite closed arms (each 50 cm in length), was used as previously described^51^. Illumination levels were set at 15–16lux in the open arms and 5–7 lux in the closed arms. Animal movement was monitored using a video-tracking system (Ethovision XT 17.5, Noldus Information Technology), which recorded locomotor trajectories, total distance travelled, and time spent in each arm. The apparatus was cleaned with a 5% ethanol solution before testing and between animals.

#### Open field test/novel object (OF/NO)

The test was conducted in a circular arena with a diameter of 100 cm. Illumination in the center of the arena was maintained at 8–10 lx. Animals were placed in the arena facing the wall and allowed to explore freely for 10 minutes. Following this initial exploration period, a novel object was positioned in the center of the arena, and animals were allowed to interact with it for an additional 5 minutes. For behavioural analysis, a virtual central zone was defined and used as an index of anxiety-like behaviour. Animal movements were recorded using a video-tracking system (Ethovision XT 17.5, Noldus Information Technology), enabling quantification of time spent in each zone and total distance travelled. The apparatus was cleaned with a 5% ethanol solution before testing and between animals.

#### Light-dark (LD) box test

An acrylic apparatus divided into two compartments (25x33 cm each) was used. One compartment was black (dark box) and kept at 20 lux while the other was white (light box) with an illumination level of 160 lux. Animals were placed in the apparatus always facing back to the door connecting the two compartments in the dark compartment and allowed to explore the apparatus for 5 minutes. Before and in-between testing, the apparatus was cleaned with a 5% EtOH solution.

### Pharmacological manipulation of plasma testosterone

#### Testosterone supplementation in HA and aged rats

Following classification into anxiety phenotypes, HA rats were segregated into two experimental groups: HA control (HA) and HA testosterone (HA-T). Animals in the HA-T group were implanted subcutaneously (s.c.) with a slow-release testosterone pellet designed to deliver physiological plasma concentrations of testosterone over a 30-day period (Belma Technologies, T-R 30). HA control animals received a placebo implant (Belma Technologies, T-R placebo). Behavioural assessments were initiated 12 days after implantation. The same experimental protocol was used for aging experiments. We took blood from Old-T rats both pre- and 30 days post-implant to confirm efficacy in old cohorts.

#### Testosterone suppression in LA rats

To pharmacologically attenuate testosterone levels in LA rats we injected a gonadotropin-releasing hormone (GnRH) receptor antagonist (Degarelix). Following classification into anxiety phenotypes, low-anxious (LA) rats received a single s.c. injection of degarelix (2 mg/kg, dissolved in saline; SML2856, Sigma-Aldrich) or an equivalent volume of saline for the LA control group. Behavioral assessments were initiated 12 days after injection.

#### Testosterone ELISA

Animals were euthanized by decapitation, and trunk blood was collected into heparinised tubes. Samples were centrifuged at 15,000 × g to obtain plasma. For hormone quantification, 6µl of plasma were processed according to the manufacturer’s instructions and analysed using commercial ELISA kits for testosterone (Enzo Life Sciences, ADI-901-065). Standard curves were generated by plotting hormone concentration on a logarithmic x-axis against optical density on a linear y-axis, followed by fitting using a four-parameter logistic regression model. Hormone concentrations in plasma samples were calculated from the resulting curves.

### Stereotaxic surgery

Animals were anesthetized with 3% isoflurane in oxygen delivered via a CombiVet animal gas anaesthesia system (Rothacher Medical, Switzerland) and maintained at 1–1.3% isoflurane in oxygen throughout the surgery. Pre-operative analgesia was delivered by subcutaneous injection of buprenorphine (0.1 mg/kg). Animals were positioned in a stereotaxic frame and placed on a heating pad to maintain body temperature. Following surgical exposure of the skull, bilateral cranial openings were drilled using a 0.5-mm burr (Komet Dental, Germany) mounted on a Tech2000 drill handpiece (Ram Products Inc., USA). Post operative analgesia was provided via paracetamol administered in the drinking water (500 mg/700 mL; Dafalgan, Bristol-Myers Squibb, Agen, France) for seven days following surgery. Behavioural experiments were initiated 4–5 weeks after surgery.

#### Androgen Receptor knock-down

HA rats received bilateral injections into the nucleus accumbens (NAc) at two rostro-caudal sites (distance from bregma: AP +1.3 mm and 2.5 mm; ML ±1.0 mm and ±1.5 mm; DV −7.0 mm). Injections were performed at a volume of 0.8µl per site using a constant infusion rate of 0.1µl/min, delivering either AAV9-GFP-U6-h-AR-shRNA (7.70 × 10¹² genome copies/mL; Vector Biolabs) or the control virus AAV2-CMV-PI-EGFP-WPRE (3 × 10¹² viral genomes/mL; Addgene). Injectors were left in place for an additional 5 min following completion of each infusion to allow diffusion and minimize backflow of virus.

#### Astrocytic Mfn2 over-expression (OE)

For Mfn2 OE, HA rats received bilateral injections into the NAc at two rostro-caudal sites (distance from bregma: AP +1.3 mm and 2.5 mm; ML ±1.0 mm and ±1.5 mm; DV −7.0 mm). Injections were performed at a volume of 0.8 µL per site using a constant infusion rate of 0.1 µL/min, delivering either AAV9-GfaABC_1_D-Mfn2-myc-T2A-GFP (6.6 × 10¹² VG/mL) or the control virus AAV9-GfaABC_1_D -GFP (1.9 × 10¹³ VG/mL). Injectors were left in place for an additional 5 min following completion of each infusion to allow diffusion and minimize backflow. Both viral vectors were generated in house at the EPFL Bertarelli Platform for Gene Therapy.

#### RNA extraction, cDNA synthesis, and qPCR

Animals were euthanized by decapitation using a guillotine, and brains were rapidly frozen in ice-cold isopentane before storage at −80 °C. The NAc was isolated for gene expression analyses by tissue punching from 200 µm-thick coronal sections prepared on a freezing cryostat. Bilateral NAc samples were collected using a 2.0 mm diameter core punch sampler (Harris Uni-Core). Tissue punches were immediately placed on dry ice and stored at −80 °C until RNA extraction. Total RNA was extracted using the RNAqueous Total RNA Isolation Kit (Ambion, AM1912) according to the manufacturer’s instructions. RNA concentration was quantified using a NanoQuant Plate in combination with a Spark microplate reader (Tecan). For complementary DNA (cDNA) synthesis, 200 ng of total RNA were reverse transcribed using qScript cDNA SuperMix (Quanta Biosciences, 95048-500). The resulting cDNA was diluted to a final concentration of 2 ng/µL. Quantitative PCR (qPCR) reactions were performed in triplicate using 1.5 µL of cDNA, 5 µL of Power SYBR Green PCR Master Mix (Thermo Fisher Scientific, 4368708), gene-specific forward and reverse primers (3.5 µL total), and 1 µL of nuclease-free water. Amplification was carried out on an ABI Prism 7900 Sequence Detection System (Applied Biosystems) under standard cycling conditions: 95 °C for 10 min, followed by 40 cycles of 95 °C for 15 s and 60 °C for 1 min. Melt curve analyses were performed at the end of each run to verify amplification specificity. Gene expression levels were calculated using the ΔΔCt method and normalized to elongation factor 1 alpha (Eef1). Primer sequences are provided in **Supplemental Table S1**.

### Evans Blue Assay for Blood-Brain-Barrier integrity

Blood–brain barrier (BBB) permeability was assessed using the Evans Blue extravasation assay as previously described^47^. Rats received a tail vein injection of Evans Blue dye (2% w/v in 0.9% saline) at a dose of 6 µl/g of body weight. Successful systemic delivery was verified by immediate blue coloration of the tail and paws. Evans Blue was allowed to circulate for 10h to permit albumin-bound dye extravasation. Animals were then deeply anesthetized with pentobarbital (10 ml/kg, Sigma Aldrich) and transcardially perfused for 5 min with ice-cold 0.1M phosphate-buffered saline (PBS) to remove intravascular dye. Animals were subsequently decapitated, and brains were rapidly extracted. The nucleus accumbens (NAc), dorsal striatum (DS), and medial prefrontal cortex (mPFC) were micro dissected as regions of interest. A peripheral control tissue (liver) was collected in parallel. All tissue samples were weighed and placed individually into 1.5ml microcentrifuge tubes. Evans Blue was extracted by incubating tissues in 500–700µl of pure dimethylformamide (DMF), with the exact volume recorded for each sample. Samples were incubated at 55 °C for 72h, followed by centrifugation at 21,000 × g for 30 min. The supernatant was collected, and Evans Blue fluorescence was quantified using a microplate reader (Tecan Spark). Dye concentrations were calculated from a serial dilution standard curve generated with Evans Blue in DMF and normalized to tissue weight.

### Electron Microscopy

Animals were euthanized by overdose of inhalation anesthesia (isoflurane) and transcardially perfused with a fixative solution containing 2.5% glutaraldehyde and 2.0% paraformaldehyde in 0.1 M phosphate buffer (pH 7.4). Brains were collected and sectioned coronally using a vibratome at a thickness of 80 µm. Sections were post-fixed with 1.5% potassium ferrocyanide and 2% osmium tetroxide, followed by staining with 1% thiocarbohydrazide and a second incubation in 2% osmium tetroxide. Samples were subsequently stained overnight with 1% uranyl acetate, rinsed in distilled water at 50 °C, and further contrasted with lead aspartate (pH 5.0) at the same temperature. Tissues were then dehydrated through a graded ethanol series and embedded in Durcupan resin, which was polymerized between glass slides at 65 °C for 24 h. Serial electron microscopy images were acquired using a block-face scanning electron microscopy approach. The coronal section containing the nucleus accumbens was isolated using a razor blade and mounted onto an aluminium stub with conductive glue. The block was trimmed with a glass knife and subsequently imaged using a scanning electron microscope (Zeiss Merlin, Zeiss NTS) equipped with an in-chamber ultramicrotome (3View, Gatan). Sequential layers of resin (50 nm thickness) were removed from the block surface, and an image was collected after each cut. Imaging was performed using an acceleration voltage of 1.7 kV, a pixel size of 6.5 nm, and a dwell time of 1 µs. Image stacks consisting of 300–400 serial sections were acquired and aligned using FIJI software (www.fiji.sc).

The distance between tight junctions was measured across 100 serial sections per sample, and the number of discontinuous tight junctions was quantified across these sections using a cutoff threshold of 50 nm, defined as the intercellular distance measured at the direct contact interface between adjacent endothelial cells.

For three-dimensional (3D) analyses, reconstructions generated in FIJI were exported to Blender (Blender.org) and quantitatively analyzed using the Neuromorph toolset. Stereological approaches were applied to quantify astrocytic end-foot coverage of blood vessels by calculating the proportion of the vascular surface area, using the basal lamina as a reference, that was enwrapped by astrocytic end-feet.

Mitochondrial size was expressed as the mean mitochondrial area across all traced mitochondria within the region of interest. Mitochondrial density was calculated as the total number of mitochondria normalized to tissue area, whereas mitochondrial coverage was determined by dividing the cumulative mitochondrial area by the total tissue area. Mitochondria–endoplasmic reticulum (ER) interactions were quantified by calculating, for each mitochondrion, the percentage of its surface area located within 50 nm of ER membranes. This metric was then averaged across all traced mitochondria within the region of interest. Neuromorph was used to quantify the surface area and volume of endothelial cells and pericytes from the reconstructed datasets.

### Nuclei isolation, library preparation and sequencing

Isolation of the intact nuclei from the NAc, followed by single nucleus library preparation and sequencing were carried as previously described^53^. Briefly, punches from the NAc from 6 hemispheres were pooled for each isolation. All solutions were supplemented with 1 mg/ml actinomycin D (Sigma, #A9415) and RNAsin (Promega, #N2111). The isolated nuclei were filtered using a 40-µm Flowmi cell strainer (Merck, #BAH136800040). Library preparation was carried out using the 10X Genomics Chromium Single Cell 3’ Library & Gel Bead Kit v3.1 following the manufacturer’s instructions. Sequencing of the libraries were performed at a Novaseq 6000 Flow Cell using read lengths of 28 nt for read1 and 90 nt for read2, at a depth of ∼70k reads/nucleus. The reads are then aligned to the rat genome and counted using the 10X Genomics custom annotation of the rat genome assembly mRatBN7.2 (Ensembl 110).

### Single nucleus transcriptomics data analysis

Seurat (v5.5.0)^91^ was used for all snRNA-seq analyses, as previously described^53^. Seurat objects for different experimental groups were initiated first and then integrated together using the SCTransform method^92^, wih the percentage of mitochondrial RNA reads used as a regression variable. RunPCA() was used with the number of dimensions that explain >95% of the variance. The resolution to use for the FindClusters() run was optimized by starting with a resolution of 0.8, and going down in 0.05 increments (until 0.2) until each discovered cluster had >20 cells, and the number of clusters are in line with the previous literature focusing on the NAc^53,93^. Differential expression analysis was carried out for each cell type using the FindMarkers() function of the Seurat package with the following parameters: logfc.threshold = 0.1, test.use = MAST.

Pathway scoring was also applied as previously published^53^.

For Gene Ontology enrichment analyses, clusterProfiler package for R (v4.20.0) was used^94^.

### Immunohistochemistry

Animals were euthanized by decapitation using a guillotine and fresh whole brains were rapidly frozen in isopentane and stored at −80 °C until further processing. Coronal cryosections (16 µm thickness) containing the NAc were prepared and mounted onto SuperFrost Plus glass slides. Sections were post-fixed in 4% paraformaldehyde, dehydrated through graded ethanol solutions, and treated with hydrogen peroxide (H₂O₂) for 10 min. Sections were then rinsed three times for 5 min each in 1× phosphate-buffered saline (PBS). Subsequently, sections were incubated for 1 h at room temperature in a blocking solution containing 0.5% Triton X-100 and 5% normal donkey serum (Sigma-Aldrich, Buchs, Switzerland) diluted in 0.1 M PBS.

Primary antibodies: anti-Aqp4 (mouse, #ab9512, 1:200), anti-Cldn5 (mouse, #35-2500, 1:200), and anti-Collagen IV (rabbit, #ab6586, 1:500) were diluted in blocking solution. Tissue sections were then incubated overnight at 4 °C with gentle agitation, followed by three washes in PBS and incubation with appropriate secondary antibodies for 2 h at room temperature. Secondary antibodies included; [goat] anti-mouse Alexa Fluor™ 488 (#A-11029), [donkey] anti-mouse Alexa Fluor™ 647 (#A-31571) and [donkey] anti-[rabbit] Alexa Fluor™ 568 (#A-10042) (ThermoFisher Scientific; 1:1000). Sections were subsequently washed three times in PBS, counterstained with DAPI (Sigma-Aldrich, #D9542) for 10 min, rinsed again in PBS, and mounted using Fluoromount-G® (SouthernBiotech, #0100-01). Slices were imaged using a Leica SP8 confocal microscope equipped with a 20×/0.75 NA air objective. Aqp4 expression was quantified as relative coverage over collagen IV by calculating the percentage of colocalization between the Aqp4 and collagen IV signals. Cldn5 levels were quantified as Cldn5 fluorescence intensity normalized to vessel area, obtained by dividing the Cldn5 channel intensity by the area occupied by the collagen IV signal, used as a proxy for vascular area. All image analyses were performed using ImageJ (NIH). At least two sections per animal were analysed and averaged to obtain a single value per hemisphere per animal for each experimental group.

### RNA in situ hybridization (ISH)

Fluorescent in situ hybridization targeting Mfn2 transcripts was performed using the RNAscope Fluorescent Multiplex 2.0 assay in combination with rat-specific RNAscope probes (Rn-Mfn2-C3, no. 505331-C3; Advanced Cell Diagnostics). Fresh whole brains were rapidly frozen in isopentane and stored at −80 °C until processing. Coronal cryosections (16 µm thickness) containing the nucleus accumbens were prepared and mounted onto SuperFrost Plus glass slides. Sections were fixed and pretreated according to the RNAscope protocol for fresh-frozen tissue. Probe hybridization was carried out using a HybEZ Hybridization System, followed by sequential signal amplification. Sections were subsequently counterstained with DAPI and coverslipped using ProLong Gold mounting medium (Thermo Fisher Scientific, #P36934). Confocal images were acquired using a Leica SP8 confocal microscope equipped with a 40×/1.25 NA glyc objective at the Bioimaging and Optics Platform (BIOP, EPFL). For experiments assessing Mfn2 expression in astrocytic populations, immunostaining for aquaporin-4 (Aqp4) was performed sequentially following the FISH procedure. Quantification of Mfn2 mRNA was restricted to Aqp4-positive regions, and results were expressed as the number of Mfn2 mRNA puncta per Aqp4-positive area. Cell detection, region segmentation, and mRNA signal quantification were performed using QuPath sOFware.

### In vitro endothelial & astrocyte co-cultures

#### Cell Culture

Primary human fetal astrocytes isolated from the cerebral cortex were purchased from ScienCell (#1800, ScienCell, USA) and cultured in Corning® BioCoat® Collagen I-coated cell culture flasks (#354485, Corning, USA) using astrocyte medium (#1801, ScienCell). Astrocytes were used at passages <5. For experiments, astrocytes were seeded at a density of 12,000 cells/cm² onto plates coated for at least 1 h with 2 µg/cm² poly-L-lysine (PLL, #P4707, Sigma-Aldrich/Merck, Germany) following washing with phosphate-buffered saline (PBS). Human Brain Microvascular Endothelial Cells (HBMVEC) were obtained from iXCells Biotechnologies (#IXC-10HU-051, iXCells Biotechnologies, USA) and cultured in Endothelial Cell Growth Medium (#MD-0010, iXCells Biotechnologies). HBMVEC were used at passages <5. For experiments, cells were seeded at a density of 20,000 cells/cm². All primary cells were maintained at 37 °C in a humidified incubator with 5% CO₂. All donors were reported as normal.

#### Drug Treatment

Testosterone (#T1500, Sigma-Aldrich/Merck) was used at concentrations of 0.1 nM, 5 nM, and 100 nM. Flutamide (#F9397, Sigma-Aldrich/Merck) was used at a concentration of 100 nM. Both compounds were dissolved in ethanol. Recombinant human TNF-α (#PHC3015L, Thermo Fisher Scientific) was used at a final concentration of 200 ng/mL

#### Cell Viability

To exclude potential cytotoxic effects of drug treatment, cell viability was assessed using the CellTiter-Glo® 2.0 Cell Viability Assay (#G9241, Promega, USA) according to the manufacturer’s instructions. Cells were seeded into white 96-well plates and treated on day 4 after seeding. Treatments were repeated 12 h later and again after 24 h. The assay was performed 48 h after the initial treatment. Luminescence was measured using a CLARIOstar Plus microplate reader (BMG LABTECH, Germany) and normalised to blank wells.

#### Transendothelial Electrical Resistance (TEER) Measurements

To assess changes in endothelial barrier properties, HBMVEC were seeded onto the apical side of Millicell® 24-well hanging cell culture inserts (#PCSP24H48, Sigma-Aldrich/Merck) and allowed to adhere for 48 h. On day 3, inserts were transferred to a cellZscope 2 (nanoAnalytics, Germany) with fresh medium and equilibrated for 24 h. Drug treatment was initiated on day 4 and repeated after 12 h and 24 h. TEER was read 48 h after the initial treatment and normalised to blank controls. For endothelial-astrocyte co-culture experiments, astrocytes were seeded onto the basal side of the inserts 24 h prior to endothelial cell seeding and proceeded as described above.

### Statistical analysis

Data analysis was performed using GraphPad Prism (v10), R (v4.3.3), and SPSS Statistics (v13.0; SPSS, Chicago, IL). The versions of individual R packages are specified in the corresponding Methods sections. Data distributions were assessed for normality (Kolmogorov-Smirnov test) prior to the application of parametric statistical tests. Differences between cumulative distributions were evaluated using the Kolmogorov–Smirnov test with α = 0.01. Comparisons between two groups were conducted using unpaired two-tailed Student’s t-tests or non-parametric equivalent Mann-Whitney tests. Comparisons involving multiple groups were performed using analysis of variance (ANOVA), followed by Tukey’s post hoc test. Statistical significance was defined as *P < 0.05, **P < 0.01, or ***P < 0.001. Unless otherwise indicated in the figure legends, data are presented as mean ± s.e.m., with individual data points shown as circles in bar graphs.

## Data availability

Code used for analyses will be made available to researchers for purposes of replication, validation, or further exploration provided that the request complies with ethical guidelines and data sharing policies. Any inquiries regarding data access should be directed to the corresponding author.

## Acknowledgements

We thank all current and former members of EPFL-LGC laboratory for valuable input and scientific discussions throughout this work. We thank the EPFL Center of PhenoGenomics (CPG) for animal care and support, and the EPFL Gene Expression Core Facility (GECF) for technical support with single-nucleus RNA sequencing.

## Funding

This work was supported by grants from the Swiss National Science Foundation (Grant Nos. 320030-232302, 310030_197 942 and 31003A_176206), the European Research Area Network Neuron (Swiss National Science Foundation (Project No. 31NE30_189061 Biostress), the Brain & Behavior Research Foundation, and intramural funding from the EPFL (all to C.S.).

Contributions

Conceptualization: C.S., H.C.A. and L.D.H. Methodology and data acquisition: H.C.A., D.H.Ü., C.D.G., M.B., E.G., F.H., J.L.S., L.S., S.A. Formal analysis: H.C.A., L.D.H., D.H.Ü., C.D.G., M.B., E.G., F.H., J.L.S., L.S., S.A. Data visualization: L.D.H., H.C.A., D.H.Ü., L.S. and S.A. Writing – original draft: C.S. and L.D.H., with input from all the authors. Funding acquisition: C.S. Supervision: C.S., C.M. and M.S.

---

**Extended Data Figure 1:**
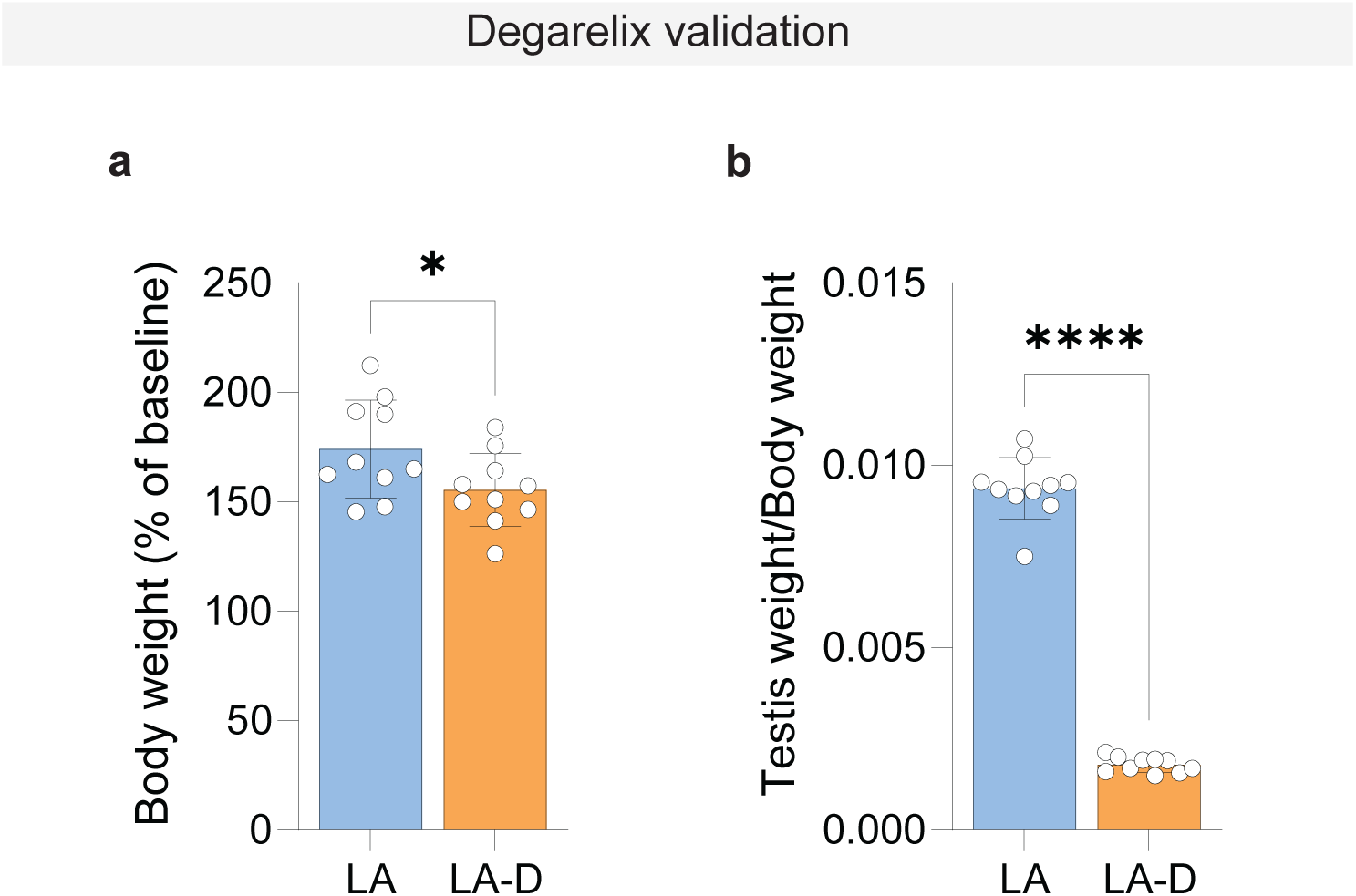
Degarelix treatment reduces body-weight gain and the testis-to-body-weight ratio in LA rats. **a,** Body weight in vehicle-treated low-anxiety (LA) and degarelix-treated low anxiety (LA-D) rats, expressed as percentage of body wait on the day of injection (LA, n=10; LA-D, n=10; unpaired t-test, *p=*0.048). **b**. Testis-to-body-weight ratio in LA and LA-D rats (LA, n=10; LA-D, n=10; unpaired t-test, *p*<0.0001). Bar plots show mean values; error bars represent s.e.m. Dots represent individual rats. *n* denotes the number of rats. ^✱^*p*<0.05 and ^✱✱✱✱^*p*<0.0001.

**Extended Data Figure 2:**
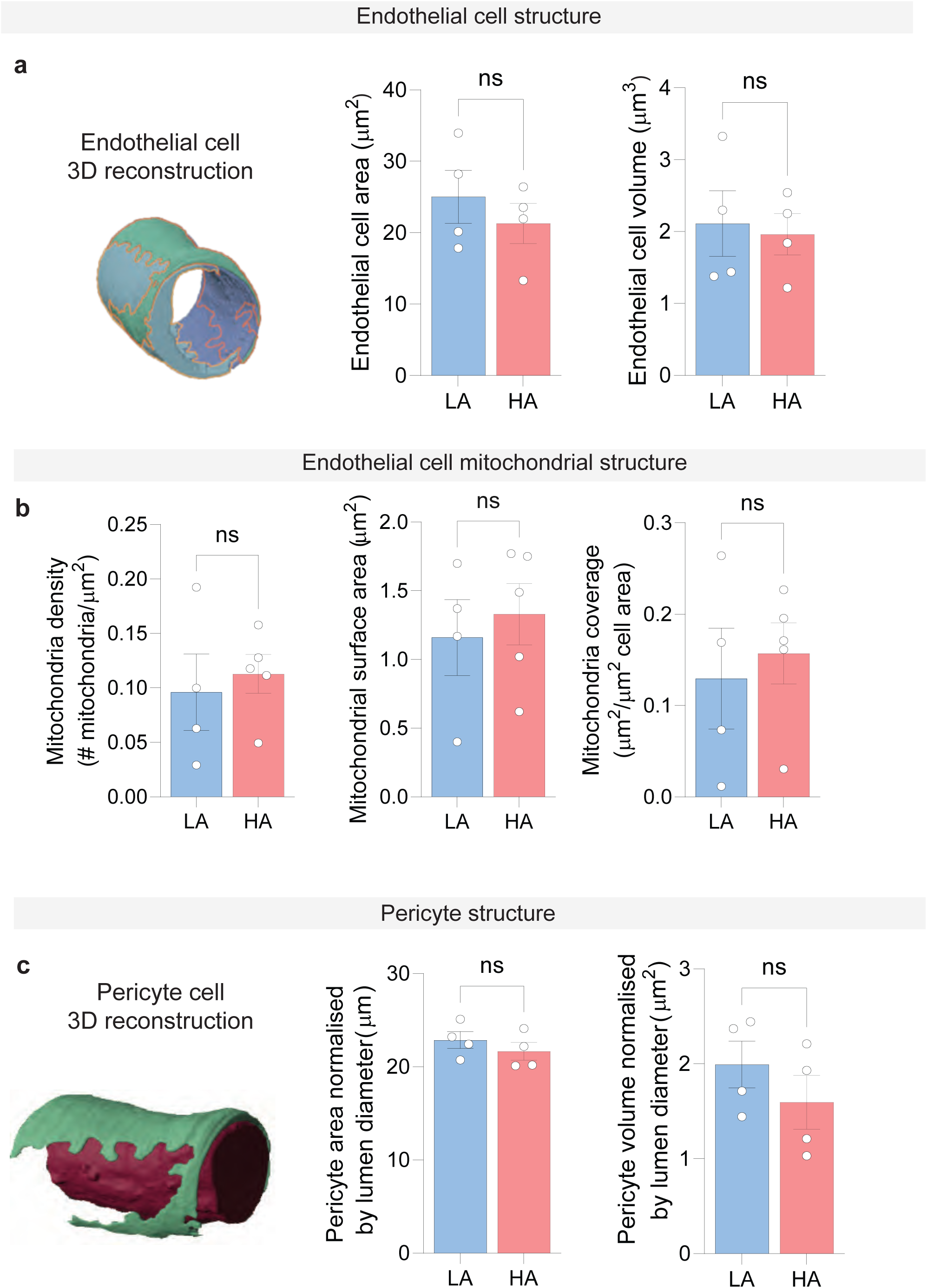
Morphology of endothelial cells and pericytes in the NAc of LA and HA rats. **a.** Representative 3D reconstruction of endothelial cells in the NAc from serial electron microscopy images and quantification of endothelial cell area (LA, n=4; HA, n=4; Mann-Whitney U test, *p*=0.686) and volume (LA, n=4; HA, n=4; Mann-Whitney U test, *p*=0.686) and volume (LA, n=4; HA, n=4; Mann-Whitney U test, *p*=0.886) of LA and HA rats **b**. Endothelial cell mitochondrial density (LA, n=4; HA, n=5; unpaired two-tailed Welch’s t-test, *p*=0.686) and volume (LA, n=4; HA, n=5; Mann-Whitney U test, *p*=0.686), mitochondrial surface area (LA, n=4; HA, n=5; Mann-Whitney U test, *p*=0.556), and mitochondrial coverage LA, n=4; HA, n=5; Mann-Whitney U test, *p*=0.730). **c**. Representative 3D reconstruction of pericytes in the NAc from serial electron microscopy images. Pericyte cell area (LA, n=4; HA, n=5; unpaired two-tailed Student’s t-test, *p*=0.390) and 3D volume (LA, n=4; HA, n=4; Mann-Whitney U test, *p*=0.343). Bar plots show mean values; error bars represent s.e.m. Dots represent individual rats. *n* denotes the number of rats. n.s., not significant.

**Extended Data Figure 3.**
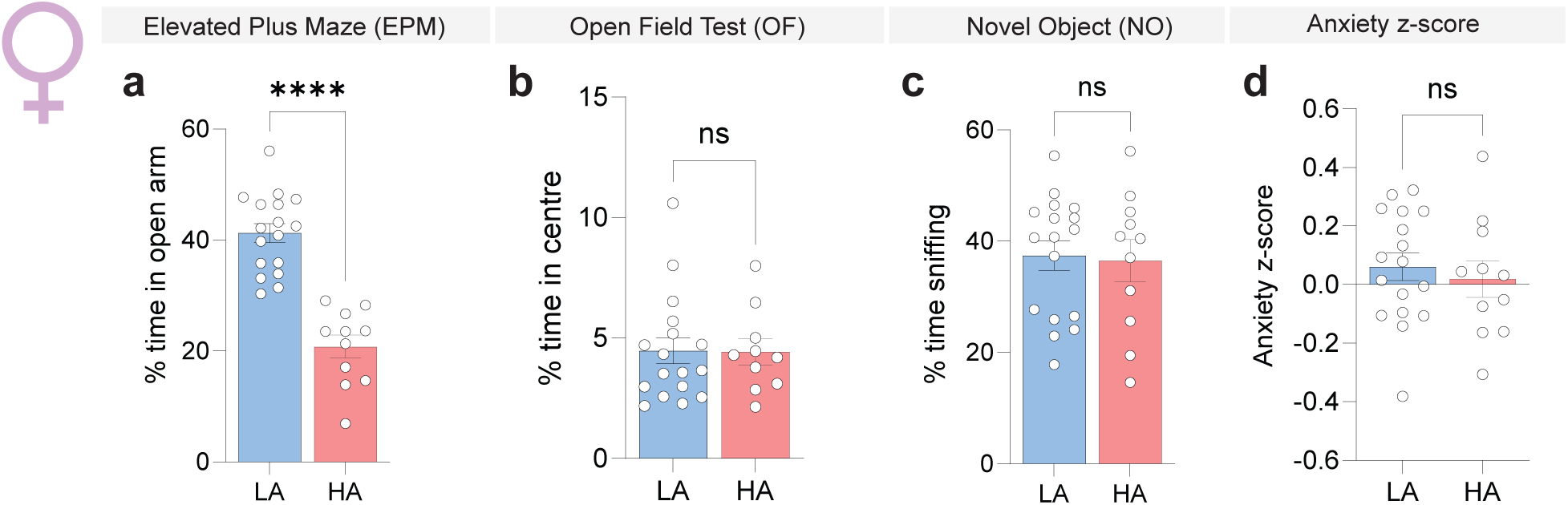
EPM-based separation of female Wistar rats does not extend across other anxiety-related measures. **a**. Percentage of time spent in the open arms of the elevated plus maze (EPM) in LA and HA female Wistar rats (LA, n=17; HA, n=11; unpaired two-tailed Student’s t-test, *p*<0.0001). **b**. Percentage of time spent in the centre of the open field test (OF) (LA, n=17; HA, n=10; unpaired two-tailed Student’s t-test, *p*=0.958). **c**. Percentage of time sniffing the novel object (N) in the NO test (LA, n=17; HA, n=10; unpaired two-tailed Student’s t-test, *p*=0.848). **d**. Composite anxiety z-score in LA and HA female rats (LA, n=17; HA, n=10; unpaired two-tailed Student’s t-test, *p=*0.596). Bar plots show mean values; error bars represent s.e.m. Dots represent individual rats. *n* denotes the number of rats. n.s., not significant, and ^✱✱✱✱^*p*<0.0001.

**Extended Data Figure 4:**
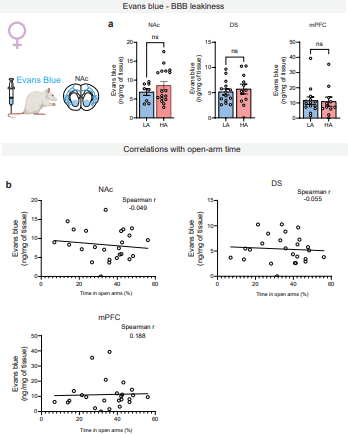
BBB permeability is not associated with anxiety-related behaviour in female Wistar rats. **a**. Evans blue accumulation in the NAc, dorsal striatum (DS) and medial prefrontal cortex (mPFC) of LA and HA female rats ( NAc: LA, n=14; HA, n=11; unpaired two-tailed Student’s t-test, *p*=0.431; DS: LA, n=17; HA, n=10; unpaired two-tailed Student’s t-test, *p*=0.611; mPFC: LA, n=17; HA, n=10; unpaired two-tailed Student’s t-test, *p*=0.867). **b**. Spearman correlations between Evans blue accumulation and percentage of time spent in the open arms of the EPM in the NAc (n = 25, r= -0.049, two-tailed *p*=0.818), DS (n = 25, r= -0.055, two-tailed *p*=0.795), and mPFC (n = 25 r= 0.188, two-tailed *p*=0.370). Bar plots show mean values; error bars represent s.e.m. Dots represent individual rats. *n* denotes the number of rats. n.s., not significant.

**Extended Data Figure 5:**
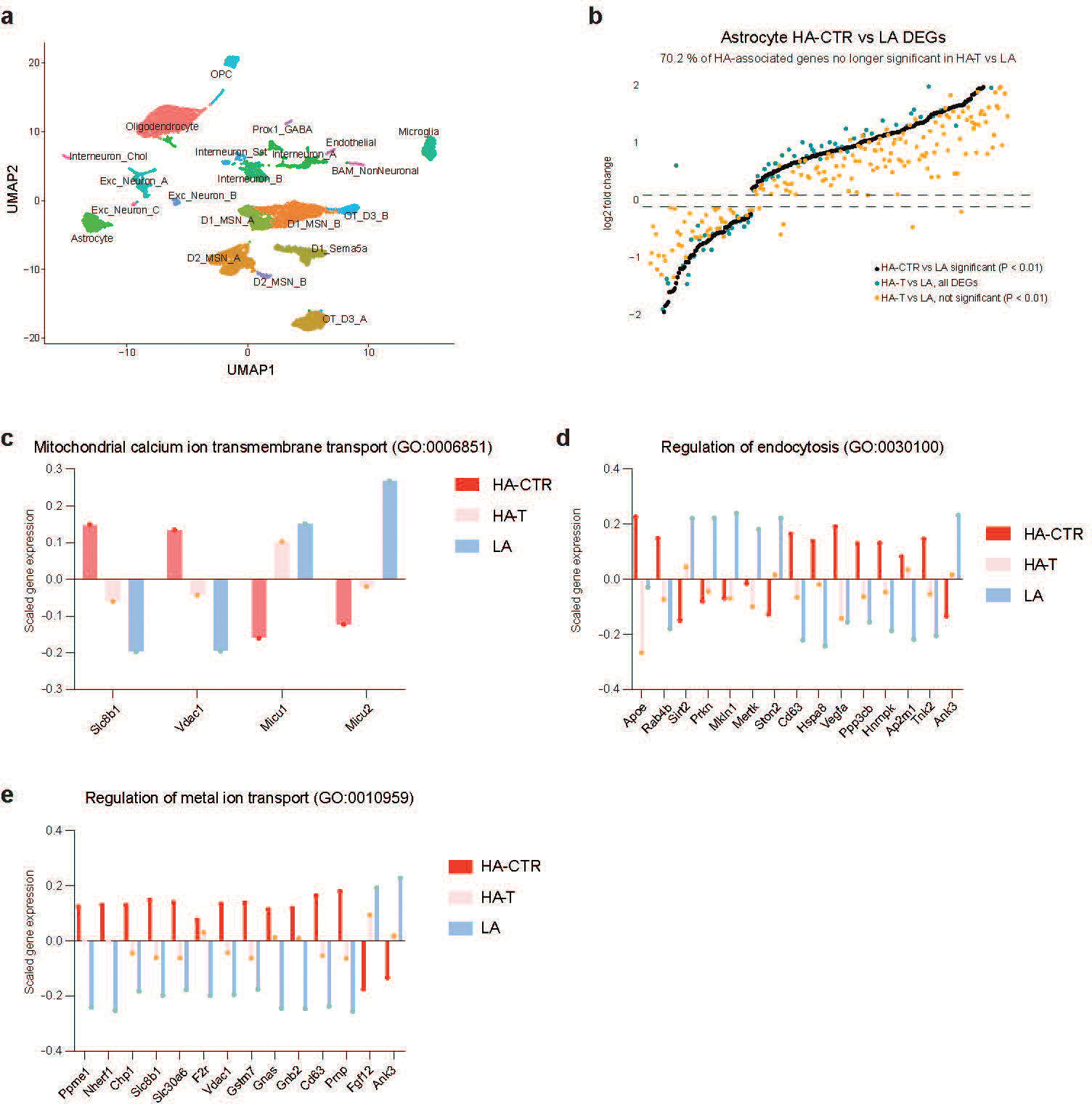
snRNA-seq analysis of testosterone-sensitive astrocytic transcriptional changes in the NAc. **a**. UMAP representation of the full NAc snRNAseq dataset, with clusters annotated by cell type. **b**. Comparison of astrocytic gene expression changes between HA-CTR, HA-T and LA rats. Genes differentially expressed between HA-CTR and LA astrocytes (*p*<0.01) are ranked by log2 fold change (black); for the same gene set, the corresponding log2 fold changes in HA-T versus LA are shown in cyan, with genes no longer significantly different between HA-T and LA (*p*>0.01) highlighted in orange. The percentage of HA-associated normalized genes no longer significantly different from LA following testosterone supplementation was calculated relative to the total number of HA-CTR versus LA differentially expressed genes. Differential expression was assessed using MAST **c-e**. Scaled expression across HA-CTR, HA-T and LA astrocytes of genes from this testosterone-shifted set contributing to selected Gene Ontology Biological Processes (GO) terms: ‘mitochondrial calcium ion transmembrane transport’ (**c**), ‘regulation of endocytosis’ (**d**) and ‘regulation of metal ion transport’ (**e**). Normalized gene counts were scaled across groups.

**Extended Data Figure 6:**
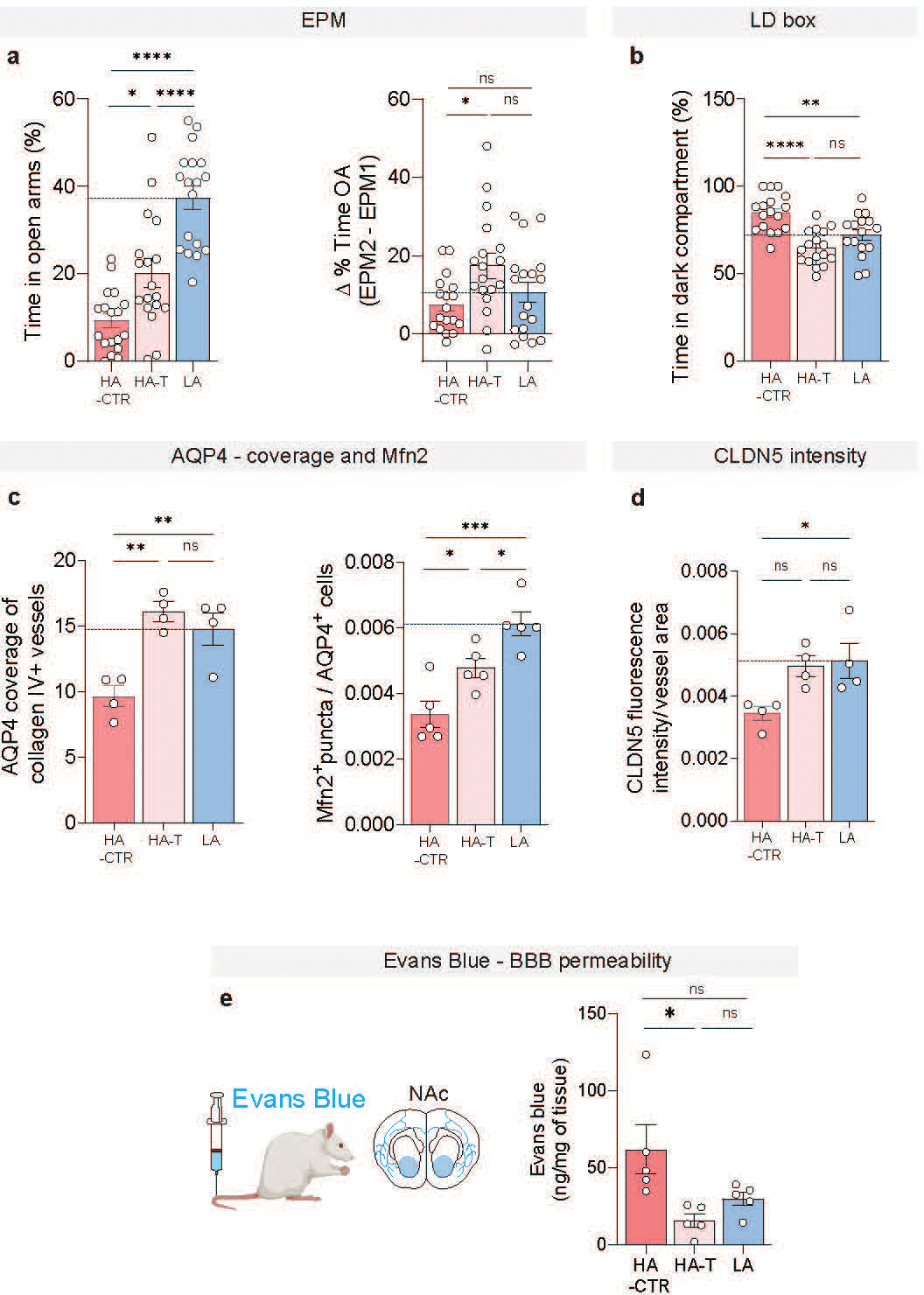
Testosterone supplementation shifts behavioural and NAc neurovascular features of HA rats towards the LA profile. **a**. Percentage of time spent in the open arms during the second elevated plus maze exposure (EPM2) in HA-CTR, HA-T and LA rats (HA-CTR, n=18; HA-T, n=18; LA. n=18; one-way ANOVA, *F*= 31.3, *p*<0.0001). Change in the percentage of time spent in the open arms between EPM1 and EPM2 (HA-CTR, n=18; HA-T, n=18; LA. n=18; one-way ANOVA, *F*= 4.25, *p*=0.020). **b**. Percentage of time spent in the dark compartment of the light-dark (LD) box in HA-CTR, HA-T and LA rats HA-CTR, n=17; HA-T, n=17; LA. n=16; one-way ANOVA, *F*= 14.6, *p*<0.0001). **c**. AQP4 coverage of Col-IV-positive vessels in the NAc of HA-CTR, HA-T and LA rats (HA-CTR, n=4; HA-T, n=4; LA. n=4; one-way ANOVA, *F*= 13.2, *p*=0.0021) and total Mfn2+ mRNA puncta per AQP4+ area (HA-CTR, n=5; HA-T, n=5; LA. n=5, One-way ANOVA: *F*= 15.3, *p*=0.0005). **d**. CLDN5 fluorescence intensity normalised to Col-IV defined vessel area in the NAc (HA-CTR, n=4; HA-T, n=4; LA. n=4; one-way ANOVA, *F*= 5.56, *p*=0.027). **e**. Evans blue accumulation in the NAc (HA-CTR, n=5; HA-T, n=5; LA. n=5; one-way ANOVA: *F*= 5.72, *p*=0.0180). Bar plots show mean values; error bars represent s.e.m. Dots represent individual rats. *n* denotes the number of rats. n.s., not significant, ^✱^*p*<0.05, ^✱✱^*p*<0.01, ^✱✱✱^*p*<0.01 and ^✱✱✱✱^*p*<0.0001.

**Extended Data Figure 7:**
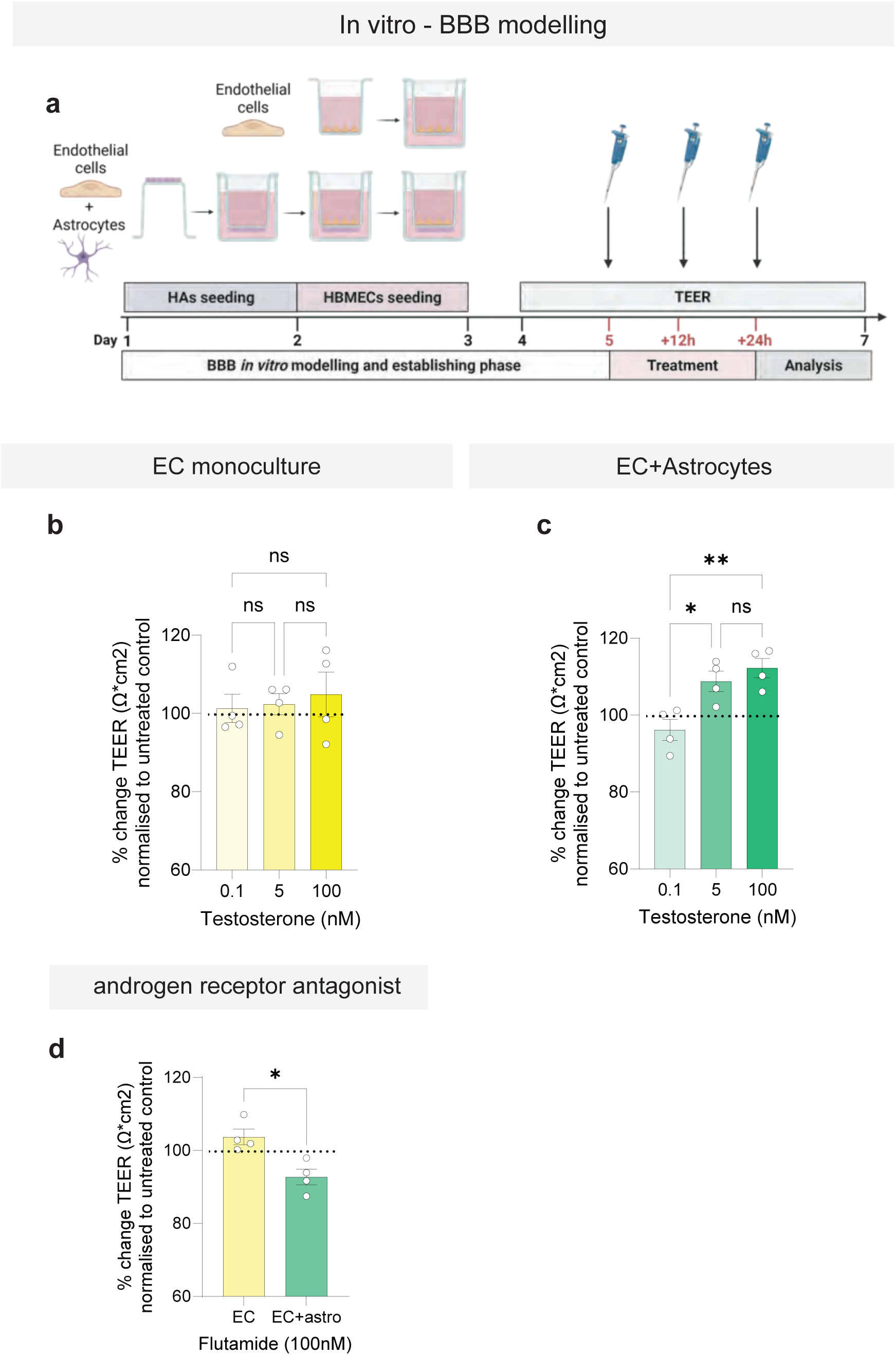
Testosterone modulates endothelial barrier resistance in an astrocyte-dependent manner *in vitro*. **a**. Schematic of the transendothelial electrical resistance (TEER) assay in human brain microvascular endothelial cell monocultures and endothelial cell-astrocyte co-cultures. **b**. TEER following treatment with testosterone (0.1, 5, and 100 nM) in endothelial cell monocultures, normalised to the corresponding untreated control (one-way ANOVA, *F*= 1.88, *p*=0.482; n=4 independent experiments). **c**. TEER following treatment with testosterone (0.1, 5, and 100 nM) in endothelial cell-astrocyte co-cultures, normalised to the corresponding untreated control (one-way ANOVA: *F*= 10.2, *p*=0.005; n=4 independent experiments, Tukey’s multiple comparisons). **d**. Effect of the androgen receptor antagonist, flutamide (100 nM) on TEER in endothelial cell monocultures (EC) and endothelial cell-astrocyte co-cultures (EC + astro), normalised to the corresponding untreated controls (EC, n=4; EC+Astro, n=4; unpaired two-tailed Student’s t-test, *p*=0.011). Bar plots show mean values; error bars represent s.e.m. Dots represent independent experiments. *n* denotes the number of cases. n.s., not significant, ^✱^*p*<0.05 and ^✱✱^*p*<0.01.

**Extended Data Figure 8:**
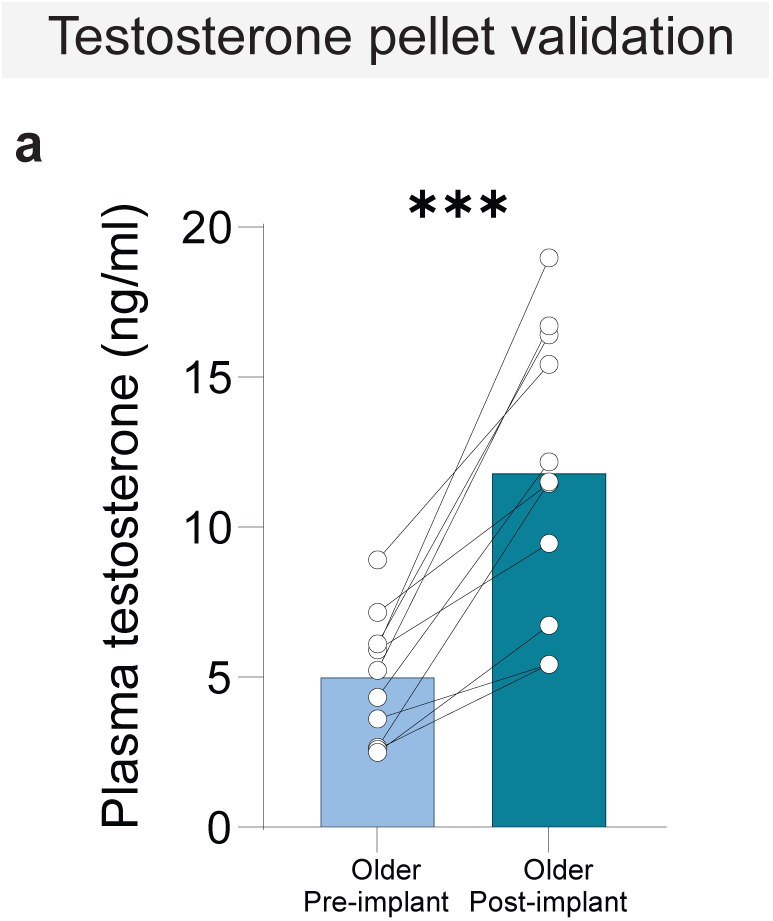
Testosterone pellet implantation increases plasma testosterone in older male rats. **a**. Plasma testosterone concentrations before and after subcutaneous testosterone pellet implantation in older male rats (n=11; paired two-tailed Student’s t-test, *p*=0.0001). Bar plots show mean values; error bars represent s.e.m. Dots represent individual rats, and connecting lines indicate paired measurements from the same animal. *n* denotes the number of rats. ^✱✱✱^*p*<0.01.

**Extended Data Figure 9:**
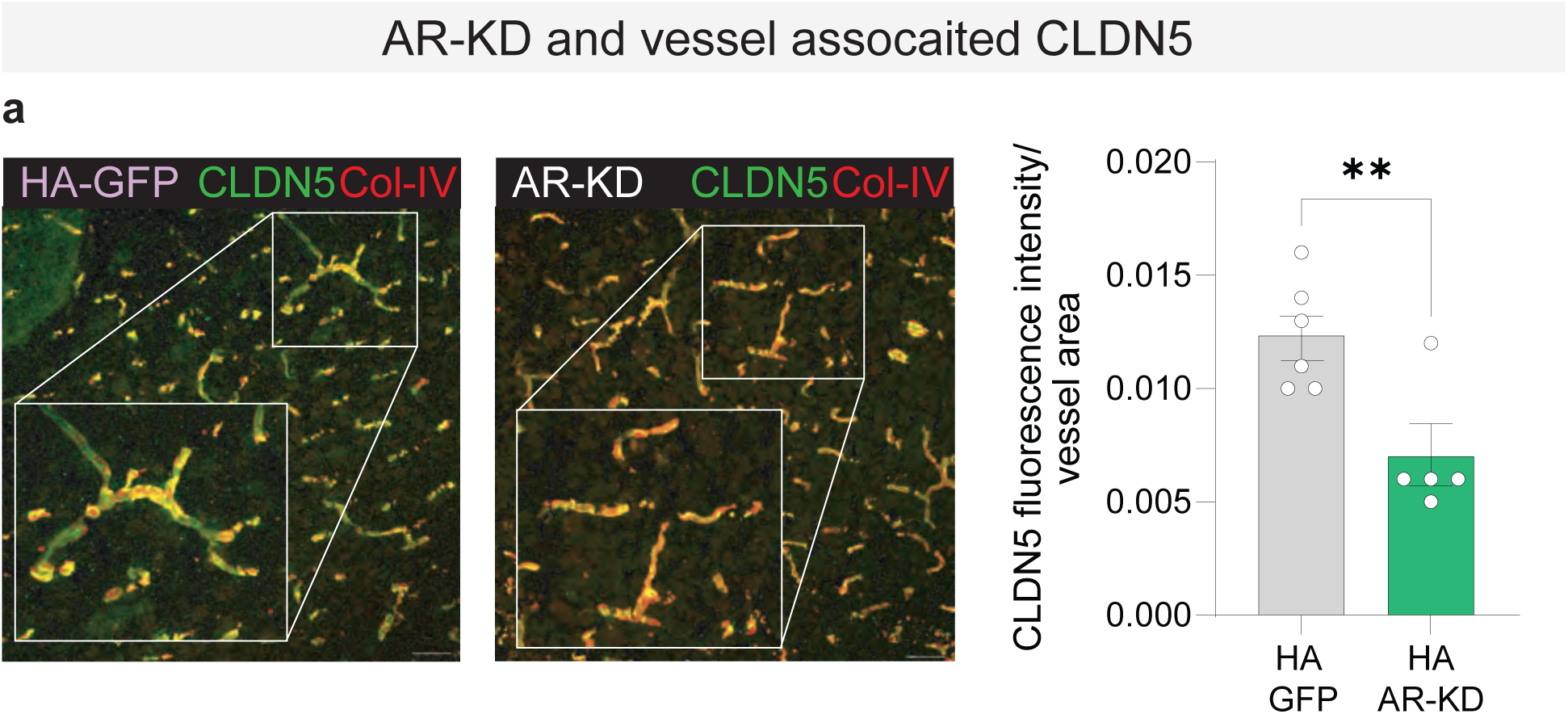
NAc androgen receptor knockdown reduces vessel-associated CLDN5 in HA rats. **a.** Representative immunofluorescence images of claudin-5 (CLDN5) and collagen IV (Col-IV) in the Nac of HA-GFP and HA AR-KD rats. Quantification of CLDN5 fluorescence intensity normalised to Col-IV defined vessel area (HA-GFP, n=6); HA AR-KD, n=5; unpaired two-tailed Student’s t-test, *p*=0.008). Scale bar = 50 µm. Bar plots show mean values; error bars represent s.e.m. Dots represent individual rats. *n* denotes the number of rats. ^✱✱^*p*<0.01.

